# Dissection of *N*-, *O*- and glycosphingolipid glycosylation changes in PaTu-S pancreatic adenocarcinoma cells upon TGF-β challenge

**DOI:** 10.1101/2021.05.14.444203

**Authors:** Jing Zhang, Zejian Zhang, Stephanie Holst, Constantin Blöchl, Katarina Madunic, Manfred Wuhrer, Peter ten Dijke, Tao Zhang

## Abstract

Pancreatic ductal adenocarcinoma (PDAC) is characterized by poor prognosis and high mortality. Transforming growth factor-β (TGF-β) plays a key role in tumor progression, which is often associated with aberrant glycosylation. How PDAC cells respond to TGF-β and the role of glycosylation therein is, however, not well known. Here, we investigated the TGF-β-mediated response and glycosylation changes in SMAD4-deficient PaTu-8955S (PaTu-S) cell line. PaTu-S cells responded to TGF-β by upregulating SMAD2 phosphorylation and target gene expression. TGF-β induced expression of the mesenchymal marker N-cadherin, but did not significantly affect epithelial marker E-cadherin expression. The differences of *N*-glycans, *O*-glycans and glycosphingolipid (GSL) glycans in PaTu-S cells with TGF-β stimulation were examined. TGF-β treatment primarily induced *N*-glycome aberrations involving elevated levels of branching, core fucosylation, and sialylation in PaTu-S cells, in line with TGF-β-induced changes in the expression of glycosylation-related genes. In addition, we observed differences in *O*- and GSL-glycosylation profiles after TGF-β treatment, including lower levels of sialylated Tn antigen, and neoexpression of globosides. Furthermore, SOX4 expression was upregulated upon TGF-β stimulation, and its depletion blocked the TGF-β-induced *N*-glycomic changes. Thus, our study provides a mechanism by which TGF-β-induced *N*-glycosylation changes in SOX4 dependent and SMAD4 independent manner in pancreatic cancer cells. Our results open up avenues to study the relevance of glycosylation in TGF-β signaling in SMAD4 inactivated PDAC.

## Introduction

Pancreatic ductal adenocarcinoma (PDAC) is one of most lethal tumors in the world as it is characterized by a poor prognosis and by a failure to respond to therapy (1). Genomic analysis of PDAC revealed that the most frequent genetic alterations include the activation of oncogene *KRAS* and inactivation of tumor suppressors tumor protein p53 (*TP53*), Sma and Mad related (*SMAD*) 4, and cyclin dependent kinase inhibitor 2A (*CDKN2A*) (2–4). The accumulation of these genetic mutations contributes to the stepwise progression of PDAC. *KRAS* mutations occurs in the early stage of PDAC (5), whereas mutations of *TP53*, *SMAD4*, and *CDKN2A* arise in advanced pancreatic intraepithelial neoplasias and invasive pancreatic adenocarcinomas (6–8). These common genetic abnormalities can profoundly perturb protein interactions and specific signaling pathways related to cell survival (9, 10), DNA damage repair (11), angiogenesis (12, 13), invasion (14), metastasis (15) and immune responses (16, 17),

The transforming growth factor-β (TGF-β) signaling pathway is involved in many cellular processes such as cell proliferation, apoptosis, migration, invasion and immune evasion, contributing to various diseases including cancer (18, 19). TGF-β is a secreted cytokine for which cellular responses are initiated by its binding to specific cell surface TGF-β type I and type II receptors, i.e., TβRI and TβRII. Upon TGF-β interaction with TβRII, TβRI is recruited and a heteromeric complex is formed (20, 21). Thereafter, the TβRII kinase phosphorylates the serine and threonine residues of TβRI, and thereby the extracellular signal is transduced across the plasma membrane (22). Subsequently, intracellular signaling by TβRI is proceeded via the phosphorylation of SMAD proteins, i.e., SMAD2 and SMAD3. Then, phosphorylated SMAD2/3 can form heteromeric complexes with the common SMAD mediator, i.e. SMAD4, which translocate into the nucleus to regulate the transcription of target genes (23). SMAD4 is a critical mediator of TGF-β-induced growth arrest (24, 25) and apoptosis (26), which results in its role as a tumor suppressor at early stages of cancer progression. However, the tumor suppressive action of TGF-β/SMAD4 signaling is lost in nearly half of PDACs because of the inactivation of SMAD4 (27). The *SMAD4* gene deletion, frameshift mutation and single point mutation leads to the deficiency of functional SMAD4 protein, which further prevents or corrupts transduction of the canonical TGF-β/SMAD4 signaling pathway. Phosphorylated TGF-β receptors can activate SMAD2/3 dependent and SMAD4 independent pathways; in that case receptor-regulated SMADs can make complexes with SRY-related HMG box 4 (SOX)4 or thyroid transcription factor-1 (TTF-1, also known as NKX2-1), which compete with SMAD4 for interaction with SMAD3 (28, 29). The SMAD3-TTF-1 and SMAD3-SOX4 complex accumulate in the nucleus, regulate the expression of pro-oncogenic TGF-β target genes and induce tumorigenesis (28–31). In addition, activated TβRI can signal via non-SMAD signaling pathways, such as the mitogen-activated protein kinase (MAPK) signaling cascade, including the extracellular signal-regulated kinase 1/2 (ERK1/2), c-Jun amino terminal kinase (JNK), p38, IκB kinase (IKK), phosphatidylinositol-3 kinase (PI3K)-AKT signaling as well as the Rho-like GTPase activity (32–36).

The SMAD-dependent and non-SMAD signaling pathways are described to be involved in multiple cellular processes including TGF-β-induced epithelial-to-mesenchymal transition (EMT) (37, 38). EMT is a crucial step towards cell metastasis, which the epithelial cells lose their polarity and cell-cell contacts and gain mesenchymal abilities such as the enhanced migratory and invasive abilities (39). EMT can be identified with morphological or phenotypic changes accompanied by a switch in the expression of EMT marker proteins; a loss of epithelial markers such as E-cadherin, claudin and an increase of mesenchymal markers including N-cadherin, Vimentin, Snail, and Slug (40, 41). EMT is a dynamic and reversible process, and cells can undergo a complete EMT or more common taken on a hybrid E/M status named “partial EMT”. A new term of epithelial– mesenchymal plasticity (EMP) is used to describe the cells undergoing intermediate E/M phenotypic states (42). The complete EMT and partial EMT both exist in PDACs, and the latter is speculated to be involved in collective cell migration and result in an enhanced metastasis rate through the formation of clusters of circulating tumor cells (43). Recent studies have provided several mechanistic bases for the TGF-β signaling in PDACs, for example, the restoration of SMAD4 in PDAC cell leads to a TGF-β-induced lethal EMT by triggering cell apoptosis via SOX4, indicating that EMT switches SOX4 function from pro-tumorigenic to pro-apoptotic (30).

Glycosylation of cellular proteins and lipids is a common posttranslational modification in cells, which affects many cellular processes such as cell adhesion, proliferation, angiogenesis, migration and invasion (44, 45). Dysregulated glycosylation is often associated with TGF-β signaling and TGF-β-induced EMT in various cancers (46), by affecting the secretion, bioavailability of TGF-β (47), TβRII localization in cells and interaction with TGF-β (48). Moreover, specific glycan structures and glycogenes involved in biosynthesis of *N*-glycans, *O*-glycans and glycosphingolipid (GSL)-linked glycans are described to increase or decrease during TGF-β-induced EMT in various cancers (49) such as lung cancer (50) and breast cancer (49, 51). During PDAC progression, aberrant glycosylation is highly correlated with several pathological processes (4, 52). Several reports have demonstrated a number of glycosylation changes, including increased fucosylation and sialylation in pancreatic cancer progression in PDACs (53, 54). The glycosylation changes of proteins such as mucin 5AC, insulin-like growth factor binding protein (IGFBP3), and galectin-3-binding protein (LGALS3BP) are involved in various signaling pathways, including TGF-β, tumor necrosis factor (TNF) and nuclear factor κ-light-chain-enhancer of activated B (NF-κ-B) pathways (55, 56). In addition, polypeptide *N*-acetylgalactosaminyltransferase 3 (GALNT3), the enzyme of the first committed step in *O*-glycosylated protein biosynthesis, can promote the growth of pancreatic cells (57). Moreover, some aberrations in glycosylation can strongly influence the properties of tumor associated extracellular matrix (ECM) and contribute to increased cell migration and invasion (58–61). Indeed, the glycan-based cancer antigen (CA)-19-9 (sialyl Lewis A) has been recognized as the hallmark for the diagnosis and early detection of pancreatic cancer (62–65). A better understanding of the aberrant glycosylation of PDAC induced by TGF-β may aid to identify potential therapeutic targets and biomarkers. However, the underlying molecular mechanisms of TGF-β signaling in SMAD4-deficient PDAC cells and their relation to glycosylation are not well understood.

In this study, the PaTu-8988S (PaTu-S) cell line, a human epithelial-like PDAC cell line with KRAS activation and inactivation of SMAD4 and CDKN2A, was employed to investigate TGF-β response and resulting glycosylation changes. Upon TGF-β treatment, PaTu-S cells were first analyzed for effects on gene expression, morphological changes, loss of epithelial traits and gain of mesenchymal markers. Next, by combining transcriptomic analysis of glycogenes with mass spectrometry glycomics, we systematically assessed the TGF-β-induced alterations in the three major classes of cell surface glycans of PaTu-S, namely, *N*-, *O*- and GSL-glycans. Furthermore, we investigated the critical role of SOX4 in TGF-β signaling and TGF-β-induced glycosylation in PaTu-S cells. This provides a stepping-stone for further studies on how glycosylation alterations contribute to TGF-β-mediated tumorigenesis of PDAC.

## Results

### TGF-β induced responses in PaTu-S cell line

To investigate whether the PaTu-S cell line that lacks SMAD4 is responsive to TGF-β treatment (Figure S1A), we performed Western blot analysis of phosphorylated SMAD2 levels of PaTu-S cells without and with TGF-β challenge. In PaTu-S cells, SMAD2 phosphorylation was significantly upregulated upon TGF-β stimulation for 1 h, which was blocked by treatment with TβRI kinase inhibitor SB431542 (Figure 1A). The expression of TGF-β target genes including cellular communication network factor 2 (*CCN2*) (66, 67), serpin family E member 1 (*SERPINE1*) (68, 69), parathyroid hormone like hormone (*PTHLH*) (70) and *SMAD7* (71) were induced by TGF-β treatment at multiple time points (Figure 1B). These genes have previously been found to be induced by TGF-β/SMAD and non-SMAD pathways and in cells in which SMAD4 was inactivated (72). In response to TGF-β stimulation for 2 days, the PaTu-S cells showed an upregulation in expression of both epithelial marker gene cadherin 1 (*CDH1*, encoding protein E-cadherin) and mesenchymal marker genes *CDH2* (encoding protein N-cadherin), SNAIL family transcriptional repressor 2 (*SNAI2*, encoding protein Slug) and *VIM* (encoding protein vimentin) (Figure 1C). At protein level, the mesenchymal marker N-cadherin was increased after TGF-β stimulation, whereas E-cadherin and Slug expression levels were not significantly affected (Figure 1D). In addition, no morphology changes of PaTu-S cells were observed after 2 days of TGF-β treatment (Figure S1B) or even longer time treatment (data not shown). The response of PaTu-S cells to TGF-β-treatment was further analyzed by immunofluorescence staining of E-cadherin and filamentous (F)-actin. We observed that TGF-β induced an increase in the formation of lamellipodia, and broadened and increased flat membrane protrusions at the leading edge of cells (Figure S1C). However, in response to TGF-β, no significant changes of E-cadherin expression and localization were observed. As an increase of lamellipodia formation has been linked to an increase in cell migration (73), we next examined the TGF-β response of PaTu-S cells using an embryonic zebrafish xenograft extravasation model (Figure S1D, E). The PaTu-S cells were pre-treated with TGF-β or vehicle control for 2 days, and thereafter a similar number of cells were injected into the circulation of zebrafish embryos. After 4 days post injection, we observed an increased number of invasive cell clusters (more than 5 cells in one cluster) in the caudal hematopoietic tissue (CHT) in the TGF-β pretreatment group compared to the non-treated group. Thus, TGF-β pre-treatment promoted the extravasation of PaTu-S cells (Figure S1D, see representative images in Figure S1E). Taken together, these results indicate that the PaTu-S cells respond to TGF-β with an upregulation of SMAD2 phosphorylation and target gene expression. In addition, upon TGF-β treatment an increase in mesenchymal marker expression was observed, but without a decrease or change in localization of epithelial markers. Moreover, TGF-β treatment of PaTu-S promoted the lamellipodia formation *in vitro* and cell extravasation *in vivo*.

**Figure 1.**
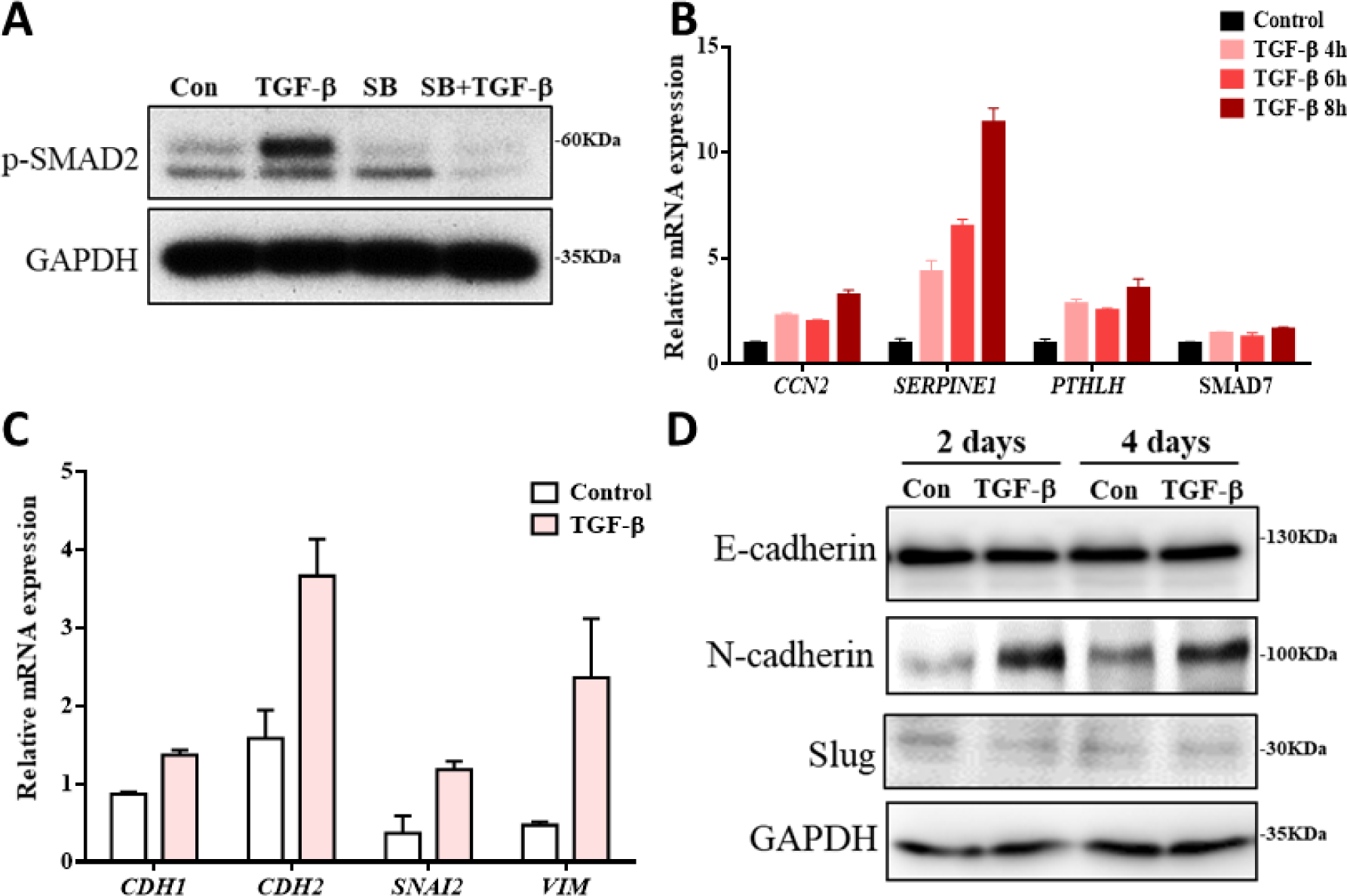
PaTu-S cell responses to TGF-β. **(A)** p-SMAD2 levels in PaTu-S cells by Western blot analysis. GAPDH, loading control. Cells were treated solely with vehicle control (Con) or TGF-β for 1 h. SB431542 (SB; 10 μM) was added for 2 h, 1 h treatment prior to TGF-β and then in combination with TGF-β for 1 h. **(B)** qRT-PCR analysis of TGF-β target genes, including *CCN2*, *SERPINE1*, *PTHLH* and *SMAD7* in PaTu-S cells treated with vehicle control or TGF-β for 4, 6 and 8 h. *GAPDH* mRNA levels were used for normalization. **(C)** qRT-PCR analysis of epithelial and mesenchymal markers including *CDH1*, *CDH2*, *SNAL2* and *VIM* in PaTu-S cells treated with vehicle control or TGF-β for 2 days. *GAPDH* mRNA levels were used for normalization. **(D)** E-cadherin, N-cadherin and Slug levels by Western blot analysis in PaTu-S cells treated with vehicle control or TGF-β for 2 days and 4 days. In latter case, fresh medium containing TGF-β or vehicle control was added after 2 days. GAPDH, loading control. Molecular weight markers are indicated on the right. Three independent experiments were performed. Representative results are shown or the data is expressed as the mean ± s.d. (n=3). TGF-β was applied at final concentration of 2.5 ng/mL.

### Differential glycosylation of PaTu-S cells with TGF-β stimulation

Changes in glycosylation of lipids and cell surface proteins have been shown to be involved in various cellular pathways, including TGF-β-mediated signaling. These pathways were found to be associated with perturbations in biological processes that contribute to cancer progression (46). After our observations regarding the effects of TGF-β on PaTu-S cells, we further investigated the differential glycosylation of PaTu-S upon TGF-β treatment. We performed a comprehensive glycomic analysis of *N*-, *O*- and GSL-glycans. Cell pellets were divided into two parts: one for the analysis of *N*- and *O*-glycans (74), and the other for the analysis of GSL-glycans. For the analysis of all three glycan classes, we used PGC nano-LC-ESI-MS/MS in negative electrospray ionization mode, enabling powerful discrimination between glycan isomers (75, 76). Glycan structures were assigned on the basis of the obtained LC-MS/MS data, and general glycobiological knowledge. Relative quantification of each glycan with and without TGF-β treatment was performed. Glycomic signatures were complemented by analyzing expression levels of glycosylation-related genes in the cells. Glycan species, traits or ratios reflecting certain biosynthetic steps were calculated to facilitate the biological interpretation of the data and to relate the MS glycomics data to transcriptomics data.

### Significant differences of *N*-glycosylation in PaTu-S cells with and without TGF-β-treatment

A total of 30 major *N*-glycan isomers spanning 27 different glycan compositions were manually identified from the investigated samples. In agreement with our previous work on *N*-glycan analysis of PaTu-S cell lines based on MALDI-TOF MS (77), the *N*-glycome data of PaTu-S was found to span four main *N*-glycan classes with major amounts of oligomannose (47.0±4.4%) and complex type (39.1±6.0%), and lesser contributions of paucimannose (9.8±0.6%) and hybrid type *N*-glycans (4.1±0.7%) (Figure 2A). More than 39% of the *N*-glycans were fucosylated, and roughly 60% were sialylated. In addition, approximately 12% of oligomannose type *N*-glycans were phosphorylated. The complex *N*-glycans were primarily diantennary (∼65%), with triantennary structures also present in significant amounts (∼15%).

**Figure 2.**
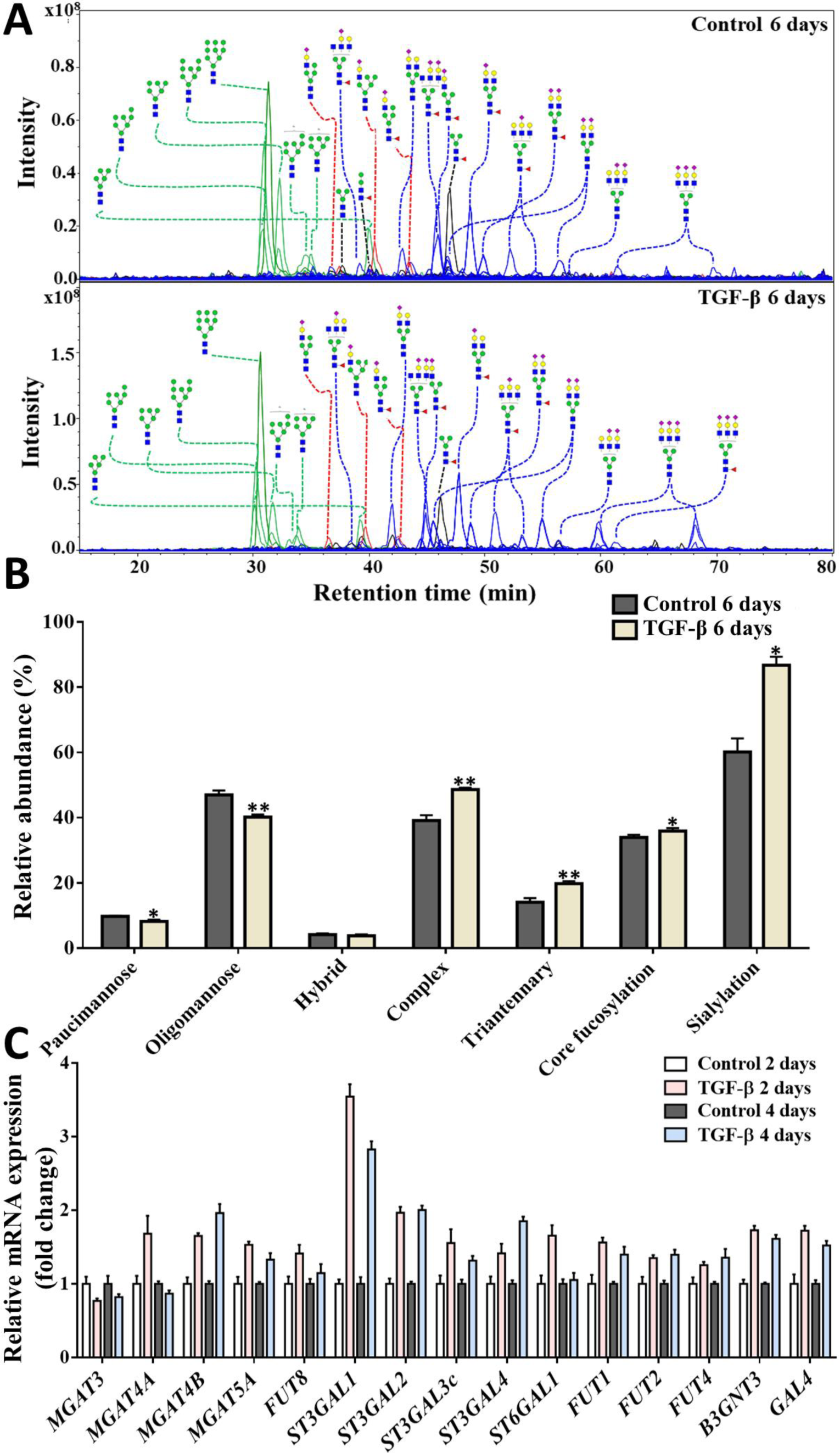
Differences of *N*-glycosylation in PaTu-S cell line with and without TGF-β treatment. **(A)** Combined extracted ion chromatograms (EICs) of *N*-glycans derived from 0.5 x 10^6^ PaTu-S cells treated with vehicle control or TGF-β for 6 days and analyzed by PGC nano-LC-ESI-MS/MS. Green trace: oligomannose; red trace: hybrid type; black trace: paucimannose; blue trace: complex type. **(B)** Relative abundance of *N*-glycan classes treated with or without TGF-β for 6 days. **(C)** qRT-PCR analysis of *N*-glycan related glycosyltransferases gene expression levels in PaTu-S cells treated with vehicle control or TGF-β for 2 days or 4 days. Gene expression levels of control groups (2 days and 4 days) were used for normalization. Representative results are shown of three independent experiments or the data is expressed as the mean ± s.d. (n=3). \**P* ≤ 0.05, \*\**P* ≤ 0.01, \*\*\**P* ≤ 0.001. Fresh medium with TGF-β (2.5 ng/mL) or vehicle control was added every 2 days in all experiments. Blue square: *N*-acetylglucosamine; green circle: mannose; yellow circle: galactose; red triangle: fucose; purple diamond: *N*-acetylneuraminic acid.

Upon TGF-β treatment, various *N*-glycosylation changes were observed. Complex *N*-glycans were increased and accounted for 49.0±1.4% of the *N*-glycome after TGF-β treatment in contrast to 39.1±6.0% in controls (Figure 2B). Specifically, triantennary *N*-glycans increased from 14.1±3.8% to 21.1±3.5% with TGF-β-treatment (Figure 2B), which is exemplified by the rather late-eluting, triantennary, trisialylated *N*-glycan species (Figure 2A). These data are in accordance with the significant upregulation of *N*-acetylglucosaminyltransferase (*MGAT*) *4A, MAGT4B* and *MGAT5A* transcripts, three genes that encode for enzymes involved in the synthesis of triantennary *N*-glycans, upon TGF-β treatment for 2 days and 4 days (Figure 2C). Notably, sialylation was high in PaTu-S cells after TGF-β treatment (0.9 sialic acid per glycan on average) with a relative abundance at 90.2±9.3%, compared to 60.1±11.2% under control conditions, which is in line with the expression patterns of β-galactoside α-2,3-sialyltransferase (*ST3GAL*)*2*, *ST3GAL3*, *ST3GAL4*, and β-galactoside α-2,6-sialyltransferase 1 (*ST6GAL1)* (Figure 2C). In addition, a slightly higher level of core fucosylation was observed (Figure 2B), in line with the upregulation of fucosyltransferase 8 (*FUT 8*) (Figure. 2C). Moreover, TGF-β treatment induced decrease of oligomannose *N*-glycans from 47.0±4.4% to 39.7±1.6% in PaTu-S cell line (Figure 2B). For a complete overview of the glycan quantification data and glycosylation-related gene expression levels see Figure S2.

### Differences of *O*-glycosylation in PaTu-S cells with and without TGF-β-treatment

Following *N*-glycan analysis, we analyzed 22 *O*-glycan species spanning 16 glycan compositions with relative quantification (Figure S3). The *O*-glycans mainly consisted of core 1 (∼52%) and core 2 structures (∼41%), with low levels of core 4 structures also present (∼2%). High level of sialylation (more than 97%) was observed in PaTu-S-derived *O*-glycans, while roughly 2% of the structures were fucosylated. The *O*-glycan profiles were very similar in TGF-β treated PaTu-S cells and non-treated cells (Figure 3A) showing a few minor and consistent changes. A significant increase of core 1 *O*-glycans was observed together with a decrease of core 2/4 *O*-glycans (Figure 3B), probably resulting from the significant upregulation of *C1GALT1* and *GCNT3* (Figure 3C). A lower level of sialyl Tn antigen was detected upon TGF-β treatment (Figure 3B), which is in line with the slightly decreased *ST6GALNAC1* (Figure 3C). For sialylation, a significant increase of α2,3 sialylation of galactose from 54.5±1.2% to 62.1±0.5% was observed (Figure 3B) in accordance with the elevated levels of *ST3GALs* (Figure 2C), and in parallel to the sialylation differences of the *N*-glycome (Figure 2B). In contrast, α2,6 sialylation of galactose, α2,6 sialylation of GalNAc, α1,2 fucosylation of galactose or α1,3/4 fucosylation stayed unaffected (Figure 3B). Interestingly, we detected no difference of α2,6 sialylation of GalNAc between PaTu-S cells with and without TGF-β stimulation (Figure 3B), despite the upregulation of α2,6 sialyltransferase related genes *ST6GALNACs (2, 3* and *4)* (Figure 3C). For a complete overview of the glycan quantification data and glycosylation-related gene expression levels see Figure S3.

**Figure 3.**
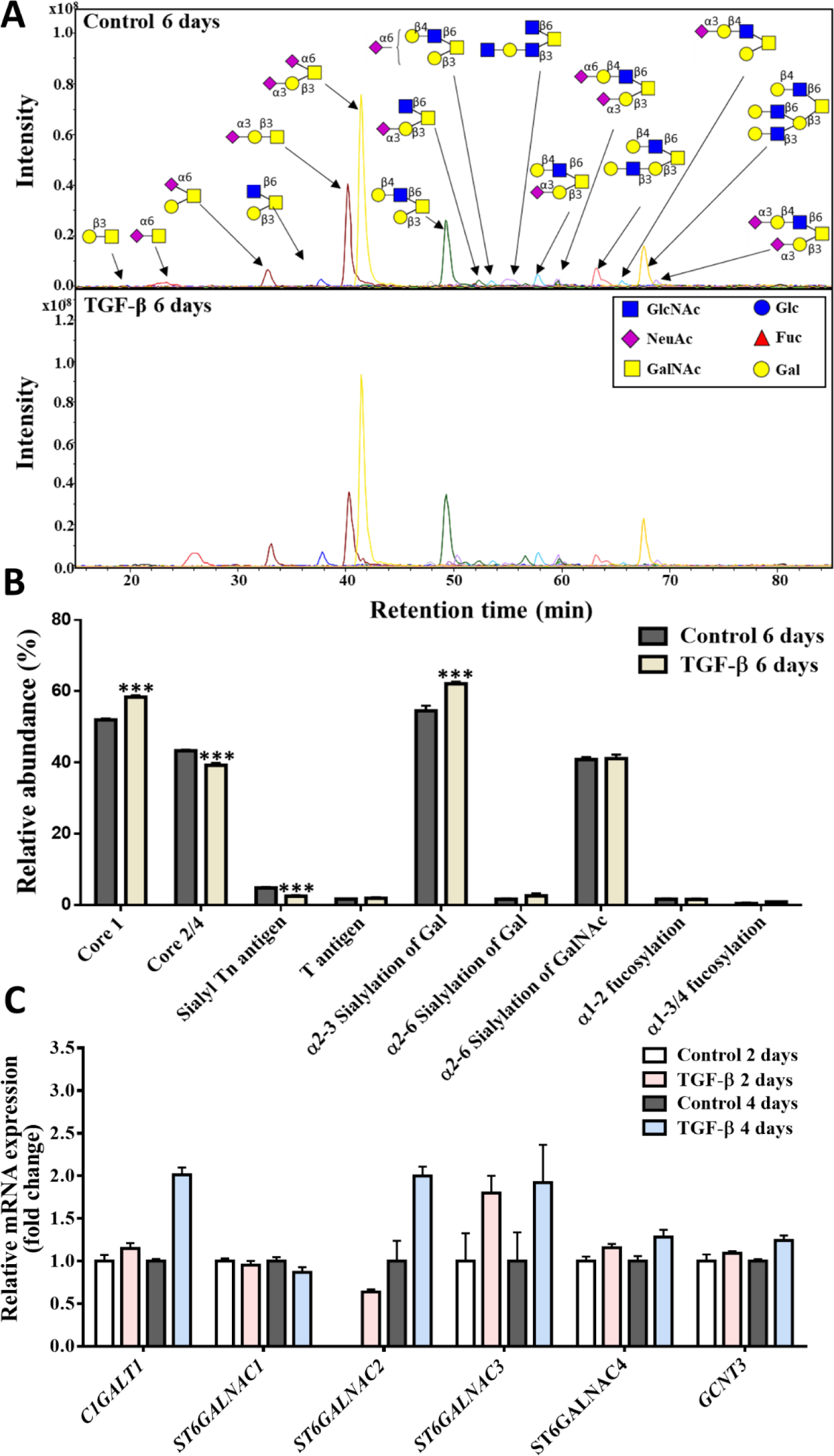
Differences of *O*-glycosylation in PaTu-S cell line without or with TGF-β treatment. **(A)** Combined EICs of *O*-glycans derived from 0.5 x 10^6^ PaTu-S cells treated with vehicle control or TGF-β for 6 days; both conditions show similar *O*-glycan profile. **(B)** Relative abundance of structural *O*-glycan classes in PaTu-S cells treated with vehicle control or TGF-β for 6 days. **(C)** qRT-PCR analysis of *O*-glycosylation related gene expression levels in PaTu-S cells treated with vehicle control or TGF-β for 2 days or 4 days. Gene expression levels of control groups (2 days and 4 days) were used for normalization. Representative results are shown of three independent experiments or the data is expressed as the mean ± s.d. (n=3). \**P* ≤ 0.05, \*\**P* ≤ 0.01, \*\*\**P* ≤ 0.001. Fresh medium containing TGF-β (2.5 ng/mL) or vehicle control was added every 2 days in all experiments.

### Neoexpression of globosides in GSL-glycomics of PaTu-S cells with TGF-β-treatment

Next, GSL-glycans were analyzed after enzymatic release of the glycan head group using EGCase I from purified GSLs derived from 2 × 10^6^ PaTu-S cells. GSL-Glycan profiles stayed largely unchanged upon TGF-β treatment except for the globoside fraction, which appeared to be specifically induced by the treatment, albeit with overall low expression levels compared to the other GSL classes (Figure 4A, 4B; Figure S4). Importantly, globosides (Gb3 and Gb4; 0.2±0.1%) were specifically present in PaTu-S after TGF-β treatment, as there was no globoside detected in the untreated sample (Figure 4B). In accordance with the glycomic data, upon TGF-β treatment, gene transcript data of day 2 also showed a 12-fold increased expression of *A4GALT*, which is the key gene for the biosynthesis of globosides (Figure 4C). Although the expression levels of multiple GSL-related genes were promoted by TGF-β stimulation for 4 days (Figure 4C), there was no significant difference in gangliosides, nsGSL, α2,6 sialic acid on galactose, α2,3 sialic acid on galactose, and fucosylation (Figure 4B). For a complete overview of the GSLs quantification data and GSL-related gene expression levels see Figure S4.

**Figure 4.**
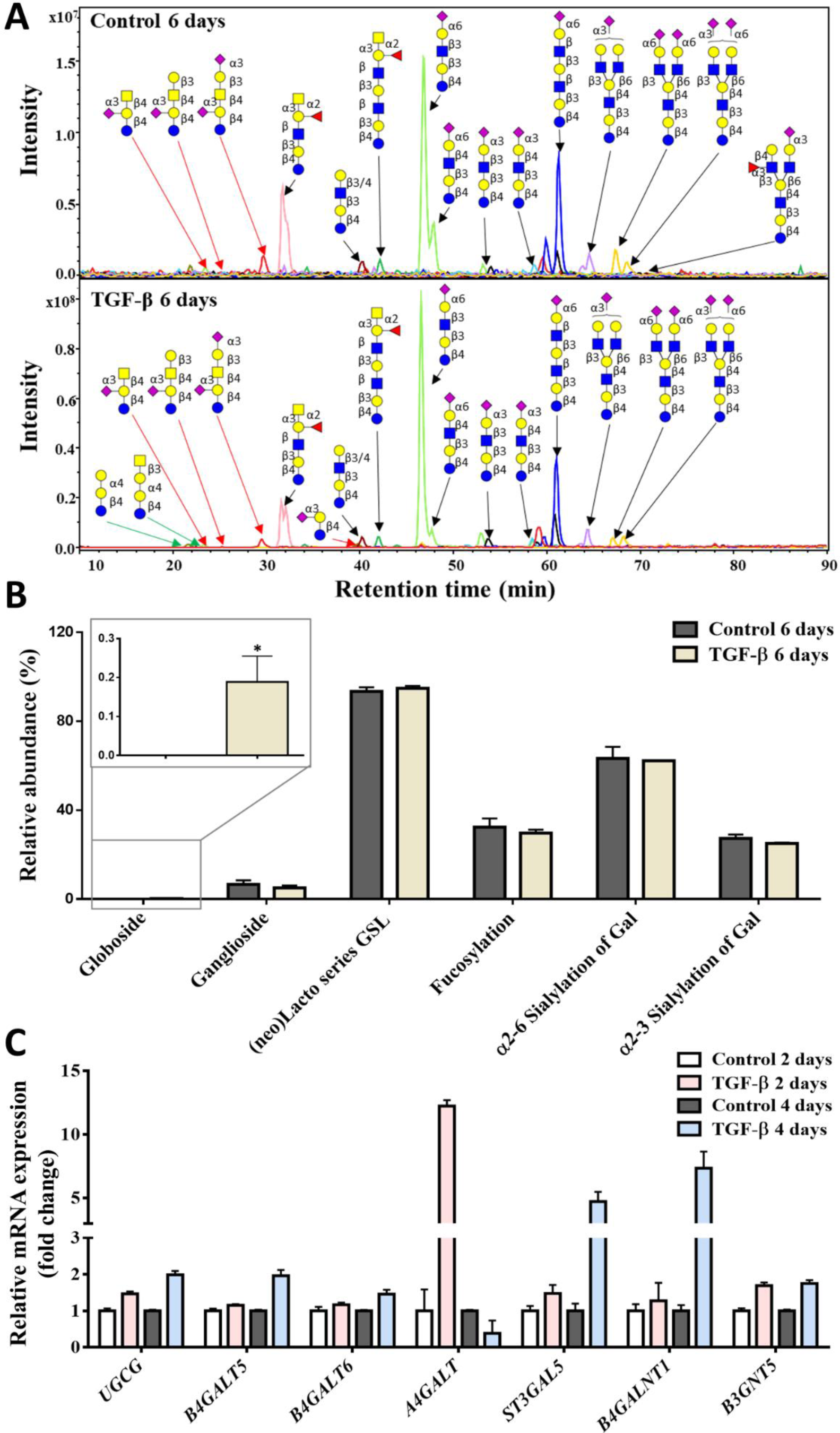
Differences of glycosphingolipid (GSL)-glycans in PaTu-S cell line without or with TGF-β treatment. **(A)** Combined EICs of GSL-glycans derived from 2 x 10^6^ PaTu-S cells treated with vehicle control or TGF-β for 6 days. Green arrow: globosides, red arrow: gangliosides, black arrow: (neo)lacto-series GSLs. **(B)** Relative abundance of structural GSL-glycan in PaTu-S cells treated with vehicle control or TGF-β for 6 days. **(C)** qRT-PCR analysis of glycosphingolipid related gene expression levels in PaTu-S cells treated with vehicle control or TGF-β for 2 days or 4 days. Gene expression levels of control groups (2 days and 4 days) were used for normalization. Representative results are shown of three independent experiments or the data is expressed as the mean ± s.d. (n=3). \**P* ≤ 0.05, \*\**P* ≤ 0.01, \*\*\**P* ≤ 0.001. Fresh medium containing TGF-β (2.5 ng/mL) or vehicle control was added every 2 days in all experiments. Blue square: *N*-acetylglucosamine; green circle: mannose; yellow circle: galactose; red triangle: fucose; purple diamond: *N*-acetylneuraminic acid.

### SOX4 is required for TGF-β-induced promotion of *N*-glycosylation

SOX4 is a key transcriptional target of TGF-β signaling pathway (28) in various cell types including breast epithelial cells (78), glioma cells (79) and pancreatic cancers (30). Importantly, SOX4 is induced by TGF-β in a SMAD4-independent manner and promotes tumorigenesis in SMAD4-null pancreatic ductal adenocarcinoma cells (30). Similarly, we found that SOX4 was upregulated in PaTu-S cells upon TGF-β stimulation for 2 days and 4 days both at mRNA (Figure 5A) and protein expression level (Figure 5B). To investigate the role of SOX4 in regulating the TGF-β-induced changes in glycan profile, we depleted the SOX4 using two independent shRNAs in PaTu-S cells. Q-PCR assay for *SOX4* expression revealed a knock down efficiency of 60% (Figure 5C). The SOX4 depletion attenuated the TGF-β-induced upregulation of *N*-glycosylation related genes including *MGAT4A* and *MGAT4B* (Figure 5D, Figure S5). Accordingly, the relative abundance of complex and specifically triantennary *N*-glycans did not increase anymore with SOX4 depletion even after TGF-β treatment for 6 days (Figure 5E). The same attenuation was observed in sialylation, as a result from the lower expression of *ST3GAL2, 3, 4,* and *ST6GAL1* in the TGF-β-treated SOX4 knockdown cells compared to the empty vector group in the presence of TGF-β (Figure 5D, Figure S5). For the core fucosylation, the relative abundance in PaTu-S has increased from 32.3±2.2% to 36.6±2.3% after TGF-β stimulation for 6 days (Figure 5E). Similarly, the level of core fucosylation in SOX4 knock down cells with TGF-β stimulation for 6 days decreased to the level of the non-treated condition, showing a reversion of the TGF-β-induced increase of core fucosylation upon SOX4 knock down (Figure 5E). All of these results indicated that SOX4 is a critical mediator for TGF-β signaling in PaTu-S cell line and plays key role in the glycosylation response, particularly in increasing sialylation, branching and core fucosylation of the *N*-glycan pool.

**Figure 5.**
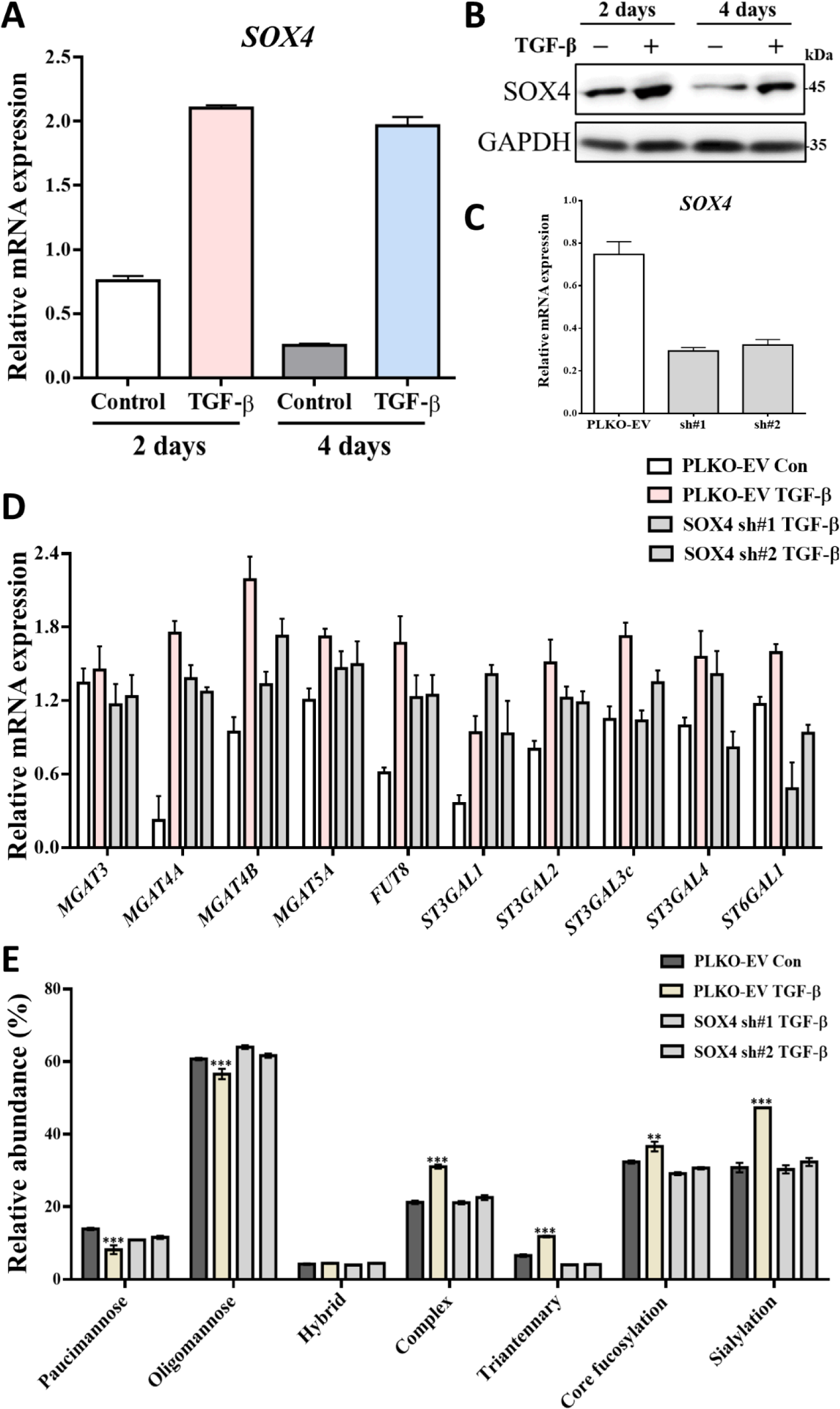
SOX4, a TGF-β target gene/protein, is needed for TGF-β-induced upregulation of *N*-glycans in PaTu-S cells. **(A)** qRT-PCR analysis of the *SOX4* in PaTu-S cells treated with vehicle control or TGF-β for 2 days and 4 days. *GAPDH* mRNA levels were used for normalization. **(B)** Immunoblotting of cell lysates for SOX4 and GAPDH (as loading control); cells were treated with vehicle control or TGF-β for 2 days and 4 days. The molecular weight markers are indicated on the right. **(C)** PaTu-S cells were stably infected with two SOX4 shRNAs (sh#1 and sh#2) or empty vector shRNA (PLKO-EV). qRT-PCR analysis was used for the expression of *SOX4* mRNA. **(D)** qRT-PCR analysis of *N*-glycosylation related genes in PaTu-S cells with PLKO-EV, SOX4 sh#1 and SOX4 sh#2 after treatment with vehicle control (Con) or TGF-β for 2 days. **(E)** Relative abundance of structural *N*-glycan classes derived from 0.5 x 10^6^ PaTu-S with PLKO-EV, SOX4 sh#1 and SOX4 sh#2 were treated with vehicle control or TGF-β for 6 days on PGC nano-LC-ESI-MS/MS in negative ion mode. Fresh medium containing TGF-β (2.5 ng/mL) or vehicle control was added every 2 days in all experiments. Representative results are shown of three independent experiments or the data is expressed as the mean ± s.d. (n=3).

## Discussion

Here, we provided a comprehensive analysis of TGF-β-induced *N*-glycan, *O*-glycan and GSL-glycan patterns in PaTu-S pancreatic adenocarcinoma cells. Our study demonstrated the significant upregulation of branching, sialylation and core fucosylation of *N*-glycans in TGF-β-treated PaTu-S cells in a SMAD4 independent manner. These *N*-glycosylation changes were found to be largely mirrored by transcriptomic changes of the underlying glycotransferase related genes. Indeed, several studies have showed that TGF-β-mediated *N*-glycosylation changes are involved in the TGF-β-SMAD4 signaling pathway but barely in SMAD4-independent pathways. For example, *ST6GAL1* and its enzymatic products including α2,6-sialylation are significantly increased during TGF-β-SMADs-induced EMT in the mouse epithelial GE11 (SMAD4 proficient) cell line, which further regulates the cell migration and invasion (80). In our study, the similar upregulation of *ST6GAL1* gene expression level and sialylation was observed in the SMAD4-deficient PaTu-S cell line. Thus, further functional studies are required to investigate the role of these certain glycosylation changes during TGF-β stimulation in SMAD4-deficient PDAC. In addition, the significant upregulation of *O*-glycosylation by TGF-β treatment, which is independent of SMAD4, in PaTu-S cell line only happened in several types of glycans such as α2,3 sialylation of galactose and core 1 structures. Importantly, the sialylated Tn antigen was notably decreased by TGF-β stimulation in Patu-S cells, which is believed to be carried by various glycoproteins and be associated with cancer progression, invasion and metastasis in some published studies (81, 82). In line with this TGF-β-induced downregulation, we have also observed the loss of sialylated Tn antigen in the mesenchymal-like Pa-Tu-8988T (PaTu-T) cells (83), which was derived from the same patient as PaTu-S cells. These results together indicate a potential role of sialylated Tn antigen in TGF-β signaling and response in PaTu-S cell lines.

GSLs were shown to be involved in the TGF-β-induced EMT process in normal murine mammary gland NMuMG (84) and human mammary carcinoma MCF7 cells (85), both cell lines express SMAD4. The promotion or inhibition of TGF-β-induced EMT and cell migration and metastasis by certain GSL, including GM2, Gg4, GM3 and GD2, have been studied in the SMAD4-dependent TGF-β response. In the SMAD4-deficient PaTu-S cell line, the only significant differences upon TGF-β stimulation were the specific expression of globosides Gb3 and Gb4. The neo-expression of globosides with TGF-β treatment is an interesting observation, since it has been reported that Gb3 is associated with tumor metastasis and invasion in many cancers, including lung cancer (86), colon cancer (87, 88), breast cancer (89), gastric adenocarcinomas (90), and pancreatic cancer (91). In our previous study, globoside Gb3 and Gb4 were specifically expressed in the mesenchymal-like PaTu-T cells compared to PaTu-S cells (83). Again, these findings indicate the globosides may contribute to the invasion and metastasis in PDAC. In the future, it will be of interest to investigate the role of sialylated Tn antigen and globosides especially Gb3 in TGF-β-promoted PDAC progression.

Importantly, we identified *SOX4*, a TGF-β transcriptional target gene, for which its gene product is required for TGF-β-induced increase of *N*-glycosylation. Knock down of SOX4 in PaTu-S cells resulted in the attenuation of TGF-β mediated upregulated branching structures, sialylation and core fucosylation. Consistent with our finding, SOX4 was shown to promote tumorigenesis in PDAC cells independent of SMAD4 expression (30). A recent investigation demonstrated that the integrin αvβ6-TGF-β-SOX4 pathway regulates multiple signaling events relevant for T cell-mediated tumor immunity in triple-negative breast cancer cells (92). Our previous study of glycosylation in PaTu-S revealed that Patu-S cells bind to dendritic cells (DCs) by galectins and other lectins, which could directly influence immune responses toward cancer cells (83). Thus, the TGF-β-SOX4 signaling pathway-induced upregulation of *N*-glycosylation especially branching, sialylation and core fucosylation might impact on immune responses.

In our study, loss of SMAD4 had no effect on the phosphorylation of upstream protein SMAD2 upon TGF-β stimulation. Interestingly, we found that TGF-β can still significantly promote the mRNA expression of TGF-β target genes including *CCN2, SERPINE1, PTHLH* and *SMAD7* in SMAD4 deficient PaTu-S cell line. This upregulation of TGF-β target genes indicates that some genes do not require SMAD4 for their regulation. Large-scale microarray analysis, which used a tetracycline-inducible small interfering RNA (siRNA) of SMAD4 (93), identified two populations of TGF-β target genes based upon their (in)dependency on SMAD4 in human immortalized keratinocytes (HaCaT) cells (72). Indeed, many TGF-β target genes such as *PTHLH* and *SMAD7* were found not to be affected by SMAD4 depletion. Although *SERPINE1* gene was classified in the category of SMAD4 dependent genes in this study (72), the authors also emphasized that this gene still displayed a residual induction with TGF-β stimulation after SMAD4 silencing (72). The previous study also demonstrated that the expression of CTGF can be induced by the combination of direct SMAD phosphorylation and indirect JAK/Stat3 activation in activated hepatic stellate cells (HSCs) (94). All these reports suggest that TGF-β target genes can be regulated by both SMAD4-dependent and SMAD4-independent pathways. Our data regarding the TGF-β-induced changes in gene expression in the SMAD4 deficient PaTu-S cell line provide further support for this notion. Moreover, we found that SOX4 knock down had no effect on TGF-β-mediated increase of target genes *CCN2, SERPINE1, PTHLH* and *SMAD7* (Figure S6). This suggests that these TGF-β-induced target genes expression are regulated in a SMAD4 and SOX4 independent manner.

We investigated the changes in expression of EMT makers upon TGF-β treatment both at gene and protein levels in PaTu-S cells. Mesenchymal markers such as N-cadherin (encoded by *CDH2* gene), Slug (encoded by *SNAL2* gene) and vimentin (encoded by *VIM* gene) are induced by TGF-β treatment for 2 days at mRNA level. At the protein level, N-cadherin (but not Slug and vimentin (data not shown) was found to be upregulated in response to TGF-β stimulation for 2 days and 4 days in PaTu-S cells. In addition, expression and localization of the epithelial marker E-cadherin are barely affected by TGF-β after 2 days and 4 days treatment. Our results indicate that PaTu-S cell line does not undergo a complete TGF-β-induced EMT. Similar results were also shown in the previous study (30) that SMAD4-mutant cells retained E-cadherin expression after TGF-β treatment for 36 h. In addition, we found that TGF-β promotes the lamellipodia formation using immunofluorescence staining of actin cytoskeleton and extravasation using zebrafish xenograft model in PaTu-S cells. Previous studies have shown that TGF-β leads to rearrangements of the actin filament system via canonical or non-canonical TGF-β signaling (95–97). In human prostate carcinoma cells, TGF-β treatment induced a rapid formation of lamellipodia through the SMAD-independent signaling pathway, which requires the activation of the Rho GTPases Cdc42 and RhoA (98). Thus, the investigation of TGF-β-induced lamellipodia formation in PaTu-S cells may offer clues of TGF-β-mediated invasion in zebrafish model and provide incentive for further studies.

In conclusion, we showed a broad-scale analysis of *N*-, O-, GSL-glycosylation changes after TGF-β treatment in PaTu-S pancreatic adenocarcinoma cells and revealed the essential role of transcriptional factor SOX4 in TGF-β-induced upregulation of *N*-glycans independent on SMAD4 expression.

## Experimental procedures

### Materials, Chemicals

Ammonium acetate, ammonium bicarbonate, cation exchange resin beads (AG50W-X8), trifluoroacetic acid (TFA), potassium hydroxide, ammonium bicarbonate, and sodium borohydride were obtained from Sigma-Aldrich (Steinheim, Germany). 8 M guanidine hydrochloride (GuHCl) was obtained from Thermo Fisher Scientific (Waltham, MA). TGF-β3 was generously provided by dr. A, Hinck (University of Pittsburg, PA). Dithiothreitol (DTT), HPLC SupraGradient acetonitrile (ACN) was obtained from Biosolve (Valkenswaard, The Netherlands) and other reagents and solvents such as chloroform, methanol, ethanol, 2-propanol, and glacial acetic acid were from Merck (Darmstadt, Germany). MultiScreen® HTS 96 multiwell plates (pore size 0.45 μm) with high protein-binding membrane (hydrophobic Immobilon-P PVDF membrane) were purchased from Millipore (Amsterdam, The Netherlands), conical 96-well Nunc plates from Thermo. The 50 mg TC18-reverse phase (RP)-cartridges were from Waters (Breda, The Netherlands). Peptide N-glycosidase F (PNGase F, lyophilized, glycerol-free) was purchased from Roche Diagnostics (Mannheim, Germany). SB431542 was from Tocris Biosciences (1614, Abingdon, United Kingdom), Alexa Fluor 488 Phalloidin was purchased from Thermo (A12379). Endoglycoceramidase I (EGCase I, recombinant clone derived from *Rhodococcus triatomea* and expressed in *Escherichia coli*) and 10x EGCase I buffer (500 mM HEPES, 1M NaCl, 20 mM DTT and 0.1% Brij 35, pH 5.2) were purchased from New England BioLabs inc. (Ipswich, MA). Ultrapure water was used for all preparations and washes, generated from a Q-Gard 2 system (Millipore, Amsterdam, The Netherlands).

### Primers and antibodies

The DNA sequences of human forward and reverse primers that were used to detect the expression of specific genes are listed in Table S1. The antibodies that were used for immunoblotting (IB) and immunofluorescence (IF): phosphor-SMAD2 1:1000 (IB: 3108, Cell signaling), GAPDH 1:1000 (IB: MAB374, Millipore), E-cadherin 1:1000 (IB, IF: 610181, BD Biosciences), N-cadherin 1:1000 (IB: 610920, BD Biosciences), Slug 1:1000 (IB: 9585, Cell Signaling), SOX4 1:1000 (IB: sc-130633, Santa Cruz), Alexa Fluor 555 secondary antibody 1:500 (IF: A-21422, Thermo).

### Cell culture

PaTu-8988S (PaTu-S) cell line was obtained from DSMZ culture bank (Braunschweig, Germany). Human embryonic kidney (HEK) 293T and A549-vimentin (VIM)-red fluorescent protein (RFP) cell lines were originally purchased from American Type Culture Collection (ATCC). PaTu-S, HEK293T and A549-VIM-RFP cells were cultured in Dulbecco’s modified Eagle medium (DMEM) with 10% fetal bovine serum (FBS) and 100U/mL penicillin-streptomycin. These cell lines were frequently tested for absence of mycoplasma contamination and authenticated by short tandem repeat (STR) profiling. For all the experiments mentioned in this study, the PaTu-S cells were always starved in DMEM with 0.5% serum for 6 h before adding the ligands.

### Western blotting

Cells were lysed with radio immune precipitation assay (RIPA) lysis buffer (150 mM NaCl, 0.1% Triton X-100, 0.5% sodium deoxycholate, 0.1% SDS, 50 mM Tris-HCl, pH 8.0, freshly added protease and phosphatase inhibitors (11836153001, Roche)) for 10 minutes (min) at 4 °C. After centrifugation at 11 × 10^3^g for 10 min at 4 °C, the protein concentrations were measured using the DC protein assay (Pierce). Equal amounts of proteins were loaded on gel, proteins separated by sodium dodecyl sulphate-polyacrylamide gel elelectrophoresis (SDS-PAGE), and thereafter protein transferred onto 45 µm Polyvinylidene difluoride (PVDF) membrane (IPVH00010, Merck Millipore). Western blotting analysis was performed by using specific primary and secondary antibodies and signals visualized with chemiluminescence. All the experiments were performed with biological triplicates and representative results are shown.

### Quantitative real-time-polymerase chain reaction (qRT-PCR)

Total RNAs were isolated using the NucleoSpin RNA II kit (740955, BIOKÉ) according to the instructions from the manufacturer. The cDNA synthesis was performed with 1 µg of RNA using RevertAid First Strand cDNA Synthesis Kit (K1621, Thermo). Real-time reverse transcription-PCR experiments were conducted with SYBR Green (Promega) in CFX Connect Detection System (1855201, Bio-Rad). Glyceraldehyde 3-phosphate dehydrogenase (*GAPDH*) mRNA levels were used to normalize specific target gene-expression. Data are shown as technical triplicates and representative of three independent biological experiments.

### Lentiviral transduction and generation of stable cell lines

SOX4 short hairpin (sh)RNAs for lentiviral transduction were obtained from Sigma (MISSION shRNA library). We tested six shRNAs and identified the most effective two shRNAs, which were then used for further experiments. These are sh#1-SOX4 (TRCN0000018217,5’-CCGGGAAGAAGGTGAAGCGCGTCTACTCGAGTAGACGCGCTTCACCTTCTTCTTTTT-3’) and sh#2-SOX4 (TRCN0000 274 207, 5’-CCGGGAAGAAGGTGAAGCGCGTCTACTCGAGTAGACGCGCTTCACCTTCTTCTTTTTG-3’). Lentiviruses were produced by transfecting HEK293T cells with SOX4 shRNA plasmids and three packaging plasmids that are pCMV-G protein of the vesicular stomatitis virus (VSVG), pMDLg-RRE (gag-pol) and pRSV-REV as described (99). The viral supernatants were harvested at 48 hours (h) post transfection and filtered through a 0.45 μm polyethersulfone (PES) filter. Virus were either directly used for infection or stored at -80 °C as soon as possible to avoid loss of titer.

To obtain stable cell lines, we first prepared a 1:1 dilution of the lentivirus in DMEM complemented with 5 ng/mL of Polybrene (Sigma). Thereafter, PaTu-S cells were infected with the lentiviral dilution at a low cell density (30%) for 24 h. After infection for 48 h, the successfully infected PaTu-S cells were selected with 2 μg/mL of puromycin for one week. The PaTu-S mCherry cells were infected with PLV-mCherry lentivirus and a single colony that highly expresses mCherry was isolated by FACS sorting, and thereafter expanded.

### Immunofluorescence staining

Cells were seeded onto sterile 18 mm-side square glass coverslips (631-1331, Menzel Gläser), complete medium was added. After overnight growth, cells were pre-starved with DMEM contained 0.5% serum for 6 h and then treated with vesicle control or with TGF-β for 2 days and 4 days. Thereafter, the cells were fixed with 4% paraformaldehyde for 30 min and permeabilized with 0.1% Triton X-100 for 10 min at room temperature. After blocking the cells with 5% bovine serum albumin (BSA) (A2058, Sigma-Aldrich) in 0.1% phosphate buffered saline (PBS)-Tween for 1 h. Subsequently, the 1:1000 diluted primary antibody of E-cadherin in PBS was added to cells for 1 h of incubation. After 3 times washing with PBS, the mixture of 1:500 diluted Alexa Fluor 555 secondary antibody and 1:1000 diluted Alexa Fluor 488 Phalloidin was added to the cells for 1 h. Thereafter, the cells were washed with PBS for 3 times and mounted with VECTASHIELD antifade mounting medium with 4′,6-diamidino-2-phenylindole (DAPI, H-1200; Vector Laboratories). Images were taken by SP8 confocal microscopy (Leica Microsystems). All the experiments were performed in biological triplicates and representative results are shown.

### Zebrafish extravasation assay

We used the transgenic green fluorescent zebrafish Tg fli:enhanced green fluorescent protein (EGFP) strain for our xenograft studies. The experiments were carried out according to the standard guidelines approved by the local Institutional Committee for Animal Welfare of the Leiden University. Zebrafish extravasation assay was performed as previous described (100). In brief, PaTu-S mCherry cells were pretreated with TGF-β for 2 days and approximately 400 cells were injected at the ducts of Cuvier into the zebrafish embryos at 48 h-post-fertilization (48-hpf). Then the injected zebrafish embryos were kept at 33°C for 4 days after injection. The latter temperature is a compromise for both the fish and cells. Thereafter, the fish were fixed with 4% paraformaldehyde and imaged by an inverted SP5 confocal microscopy (Leica Microsystems). The number of invasive cell clusters (more than 5 cells were defined as a cluster) between the vessels in the caudal hematopoietic tissue region were counted.

### Preparation of *N*- and *O*-glycan alditols released from PaTu-S cells

*N*-glycan and *O*-glycan alditols released from PaTu-S cells were prepared using a 96-well plate sample preparation method performed as previously described (74, 83). In brief, lysates from 0.5 x 10^6^ cells were applied to the hydrophobic Immobilon-P PVDF membrane in a 96-well plate format. Protein denaturation was achieved by applying 75 µL denaturation mix (72.5 µL 8 M GuHCl) and 2.5 µL 200 mM DTT) in each well, followed by shaking for 15 min and incubating at 60°C in a moisture box for 30 min. Subsequently the unbound material was removed by centrifugation.

The *N*-glycan was released by adding peptide:N-glycosidase F (PNGase F) (2 U of enzyme diluted with water to 15 µL) to each well and incubated overnight at 37°C. Released *N*-glycans were collected from the PVDF plate by centrifugation, and the glycosylamine versions of the released *N*-glycans were hydrolyzed by adding 20 µL of 100 mM ammonium acetate (pH 5), incubated at room temperature (RT) for 1 h, and dried in a SpeedVac concentrator 5301 (Eppendorf, Hamburg, Germany) at 35°C. Collected *N*-glycans were then reduced and desalted followed by PGC cleanup using a 96-well plate based protocol (74, 76). Samples were dried in a SpeedVac concentrator directly in PCR plates and re-dissolved in 10 μL of water prior to porous graphitized carbon nano-liquid chromatography (PGC nano-LC-ESI-MS/MS) analysis.

After removal of *N*-glycans, the *O*-glycans were released from the same PVDF membrane-immobilized sample via reductive β-elimination. Briefly, 50 μL of 0.5 M NaBH4 in 50 mM KOH was applied onto each PVDF membrane well after rewetting with 3 μL of methanol. Plates were placed for 15 min on a horizontal shaker and incubated in a humidified plastic box for 16 h at 50°C. After cooling to RT, released *O*-glycans were recovered by centrifugation at 1000 *g* for 2 min into 96 well collection plates. The wells were rewetted by 3 μL of methanol and washed three times with 50 μL of water with 10 min incubation steps on a horizontal shaker prior to centrifugation at 500 *g* for 2 min. Prior to desalting, the collected samples were concentrated to approximately 30 μL under vacuum in a SpeedVac concentrator at 35°C for 2 h. Subsequently, 3 μL of glacial acetic acid was added to quench the reaction followed by brief centrifugation to collect the samples at the bottom of the wells. The following high throughput desalting and porous graphitized carbon (PGC) solid phase extraction (SPE) purification were performed as described in the *N*-glycan preparation section under 2.4. The purified *O*-glycan alditols were re-suspended in 10 μL of water prior to PGC nano-LC-ESI-MS/MS analysis.

### Preparation of GSL-glycan alditols released from PaTu-S cells

Extraction of GSLs and preparation of GSL-glycan alditols from PaTu-S cells were performed in triplicate as previously described (83). Shortly, 2 x 10^6^ cells were harvested, washed and resuspended by 200 μL of water. The cell samples were lysed by vortexing and sonication for 30 min. Chloroform (550 μL) was added to the samples followed by 15 min sonication. Methanol (350 μL) was added to the cell pellets and incubated for 4 h with shaking at room temperature. The upper phase containing GSLs was collected after centrifugation at 2700 × g for 20 min. Then, 400 μL of chloroform/methanol (2:1, v/v) was added, followed by adding 400 μL of methanol/water (1:1, v/v). After sonication and centrifugation, the upper phase was collected and pooled to the previous sample. The process of adding methanol/water (1:1, v/v), sonication, centrifugation and removing upper phase was repeated for another two times. In each replicate, the upper phase was collected and replaced by the same volume of methanol/water (1:1, v/v). The combined upper phases were dried under vacuum in an Eppendorf Concentrator 5301 (Eppendorf, Hamburg, Germany) at 30°C.

Before the purification of the GSLs using reverse phase (RP) SPE, the samples were dissolved in 100 μL methanol and vortexed for 10 min, followed by the addition of 100 μL water. TC18-RP-catridges were prewashed with 2 mL of chloroform/methanol (2:1, v/v), 2 mL of methanol followed by equilibration with 2 mL methanol/water (1:1, v/v). The extracted GSLs were loaded to the cartridge 3 times and washed with 2 mL methanol/water (1:1, v/v). The GSLs were eluted from the column with 2 mL methanol and 2 mL chloroform/methanol (2:1, v/v). The samples containing the eluate were evaporated under nitrogen for 1 h and dried under vacuum in an Eppendorf Concentrator at 30°C.

To release the glycans from the GSLs, a mixture of EGCase I (12 mU, 2 μL), EGCase I buffer (4 μL) and water (34 μL) (pH 5.2) was added to each sample and incubated for 36 h at 37°C. The released glycans were collected and loaded on TC18-RP-cartridges, which had been preconditioned with 2 mL of methanol and 2 mL of water. The samples were washed with 200 ul of water and residual glycans were loaded to the cartridge. Then, 500 μL of water was added to the cartridge to wash the glycans from the column. The flow-through and wash fractions were pooled and dried in an Eppendorf Concentrator at 30°C. The reduction was carried out with slight modifications following the same procedure as described in previous work (76, 83). In brief, GSL-glycans were reduced to alditols in 20 μL of sodium borohydride (500 mM) in potassium hydroxide (50 mM) for 2 h at 50°C. Subsequently, 2 μL of glacial acetic acid was added to acidify the solution and quench the reaction. The desalting of GSL-glycans were performed as previously described. Glycan alditols were eluted with 50 μL of water twice. The combined flow-through and eluate were pooled and dried under vacuum in an Eppendorf Concentrator at 30°C. The carbon SPE clean-up was performed and the purified glycan alditols were re-suspended in 20 μL of water prior to PGC nano-LC-ESI-MS/MS analysis.

### Analysis of *N*-, *O*- and GSL-glycan alditols released from PaTu-S cells using PGC nano-LC-ESI-MS/MS

The analysis of glycan alditols was performed using PGC nano-LC-ESI-MS/MS following a method described previously (74, 83). Measurements were performed on an Ultimate 3000 UHPLC system (Thermo) equipped with a home-packed PGC trap column (5 μm Hypercarb, 320 μm x 30 mm) and a home-packed PGC nano-column (3 μm Hypercarb 100 μm x 150 mm) coupled to an amaZon ETD speed ion trap (Bruker, Bremen, Germany). Mobile phase A consisted of 10 mM ABC, while mobile phase B was 60% (v/v) acetonitrile/10 mM ABC. The trap column was packed with 5 μm particle size PGC stationary phase from Hypercarb PGC analytical column (size 100 × 4.6 mm, 5 μm particle size, Thermo) while the PGC nano-column were packed with 3 μm particle size PGC stationary phase from Hypercarb PGC analytical column (size 100 × 4.6 mm, 5 μm particle size, Thermo).

To analyze glycans, 2 μL injections were performed and trapping were achieved on the trap column using a 6 μL/min loading flow in 1% buffer B for 5 min. Separation was achieved with a multi-step gradient of B: 1-9% in 1 min and 9-49% in 80 min for *N*-glycan and 1-52% over 72 min for *O*-glycans followed by a 10 min wash step using 95% of B at a flow of rate of 0.6 μL/min. To separate GSL-glycans, a linear gradient from 1% to 50% buffer B over 73 min was applied at a 0.6 μL/min flow rate. The column was held at a constant temperature of 45°C.

Ionization was achieved using the nanoBooster source (Bruker) with a capillary voltage of 1000 V applied and a dry gas temperature of 280°C at 5 L/min and isopropanol enriched nitrogen at 3 psi. MS spectra were acquired within an *m/z* range of 500-1850 for *N*-glycans, 380-1850 for *O*-glycans, and 340-1850 for GSL-glycans in enhanced mode using negative ion mode, smart parameter setting (SPS) was set to *m/z* 1200, 900 and 900, respectively. MS/MS spectra were recorded using the top 3 highest intensity peaks.

Structures of detected glycans were studied by MS/MS in negative mode (101). Glycan structures were assigned on the basis of the known MS/MS fragmentation patterns in negative-ion mode (75,102,103), elution order, and general glycobiological knowledge, with help of Glycoworkbench (104) and Glycomod (105) software. Relative quantification of individual glycans was performed by normalizing the total peak area of all glycans within one sample to 100%. Relative abundances of specific glycan derived traits were displayed by summing relative intensities of each glycan structure containing the epitope multiplied by the number of epitopes per glycan. Structures are depicted according to the Consortium of Functional Glycomics (CFG).

### Statistical analysis

Statistical analyses were performed with a Student’s unpaired t test using Prism 8 software (GraphPad La Jolla, CA). The numerical data from triplicates are expressed as the mean ± s.d, and the results of Zebrafish assay and PGC nano-LC-ESI-MS/MS are expressed as the mean ± SEM. P value is indicated by asterisks in the figures, \**P* ≤ 0.05, \*\**P* ≤ 0.01, \*\*\**P* ≤ 0.001. *P* ≤ 0.05 was considered statistically significant.

## Acknowledgments

We acknowledge the support of the Chinese Scholarship Council (CSC) to Jing Zhang, Cancer Genomics Centre Netherlands (CGC. NL) to Peter ten Dijke and Austrian Science Fund (FWF, W1213) to Constantin Blöchl. We thank Irma van Die and Ana I. Belo for the contribution of PaTu-S cell line and fruitful discussions.

## Author contributions

Project ideation: S. Holst, P. ten Dijke Conception and design: P. ten Dijke, J. Zhang, T. Zhang, M. Wuhrer Development of methodology: J. Zhang, T. Zhang, S. Holst, K. Madunic Acquisition of data: J. Zhang, T. Zhang, Z. Zhang Analysis and interpretation of data: J. Zhang, T. Zhang, C. Blöchl Writing, review and/or revision of the manuscript: J. Zhang, T. Zhang, P. ten Dijke, M. Wuhrer Study supervision: P. ten Dijke, T. Zhang, M. Wuhrer

## Data availability

The raw mass spectrometric data files that support the findings of this study are available in GlycoPOST in mzXML format, with the identifier GPST000180, accessible via the following link https://glycopost.glycosmos.org/entry/GPST000180.

## Abbreviations

PDAC: pancreatic ductal adenocarcinoma
TGF-β: Transforming growth factor-β
GSL: glycosphingolipid
TP53: tumor suppressors tumor protein p53
SMAD: Sma and Mad related
CDKN2A: cyclin dependent kinase inhibitor 2A
TβRII: TGF-β type II receptor
TβRI: TGF-β type I receptor
MAPK: mitogen-activated protein kinase
ERK: extracellular signal-regulated kinase
JNK: c-Jun amino terminal kinase
IKK: IκB kinase
PI3K: phosphatidylinositol-3 kinase
EMT: epithelial-to-mesenchymal transition
EMP: epithelial–mesenchymal plasticity
SOX4: SRY-related HMG box 4
TTF-1: thyroid transcription factor-1
IGFBP3: insulin-like growth factor binding protein
LGALS3BP: galectin-3-binding protein
TNF: tumor necrosis factor
NF-κ-B: nuclear factor κ-light-chain-enhancer of activated B
GALNT3: polypeptide N-acetylgalactosaminyltransferase 3
ECM: extracelular matrix
CA: cancer antigen
qRT-PCR: quantitative real-time-polymerase chain reaction
PGC: nano-LC-ESI-MS/MS porous graphitized carbon nano-liquid chromatography coupled to a tandem mass spectrometer
TFA: trifluoroacetic acid
ACN: acetonitrile
RP: reverse phase
PNGase F: Peptide N-glycosidase F
EGCase I: Endoglycoceramidase I
IB: immunoblotting
IF: immunofluorescence
HEK: human embryonic kidney
VIM: vimentin
RFP: red fluorescent protein
ATCC: American Type Culture Collection
DMEM: Dulbecco’s modified Eagle medium
FBS: fetal bovine serum
STR: short tandem repeat
RIPA: radio immune precipitation assay
Min: minute
SDS-PAGE: sodium dodecyl sulphate-polyacrylamide gel elelectrophoresis
PVDF: polyvinylidene difluoride
GAPDH: glyceraldehyde 3-phosphate dehydrogenase
shRNA: short hairpin RNA
VSVG: vesicular stomatitis virus
h: hour
PES: polyethersulfone
BSA: bovine serum albumin
PBS: phosphate buffered saline
DAPI: 4’,6-diamidino-2-phenylindole
hpf: hour-post-fertilization
RT: room temperature
SPS: smart parameter setting
CFG: Consortium of Functional Glycomics
CCN2: cellular communication network factor 2
SERPINE1: serpin family E member 1
PTHLH: parathyroid hormone like hormone
CDH: gene cadherin
SNAI2: snail family transcriptional repressor 2
F-actin: filamentous actin
CHT: caudal hematopoietic tissue
MAGT: *N*-acetylglucosaminyltransferase
FUT: fucosyltransferase
ST3GAL: β-galactoside alpha-2,3-sialyltransferase
ST6GAL1: β-galactoside alpha-2,6-sialyltransferase 1
CRC: colorectal cancer
DCs: dendritic cells
HaCaT: human immortalized keratinocytes
HSCs: hepatic stellate cells
siRNA: small interfering RNA
EICs: extracted ion chromatograms

