## Supplemental materials for "Dissection of *N*-, *O*- and glycosphingolipid glycosylation changes in PaTu-S pancreatic adenocarcinoma cells upon TGF-β challenge"

**Supporting Information**

Figure S1. Responses to TGF-β in PaTu-S cell line.

Figure S2. Changes of *N*-glycosylation in PaTu-S cell line without or with TGF-β treatment.

Figure S3. Changes of *O*-glycosylation in PaTu-S cell line without or with TGF-β treatment.

Figure S4. Changes of glycosphingolipids (GSLs) in PaTu-S cell line without or with TGF-β treatment.

Figure S5. mRNA expression levels of *N*-glycan biosynthesis related glycosyltransferases genes in PaTu-S cells with SOX4 deletion.

Figure S6. mRNA expression levels of TGF-β target genes in PaTu-S cells with SOX4 deletion.

Table S1. Primers used in this study for quantitative real-time-polymerase chain reaction (qRT-PCR)

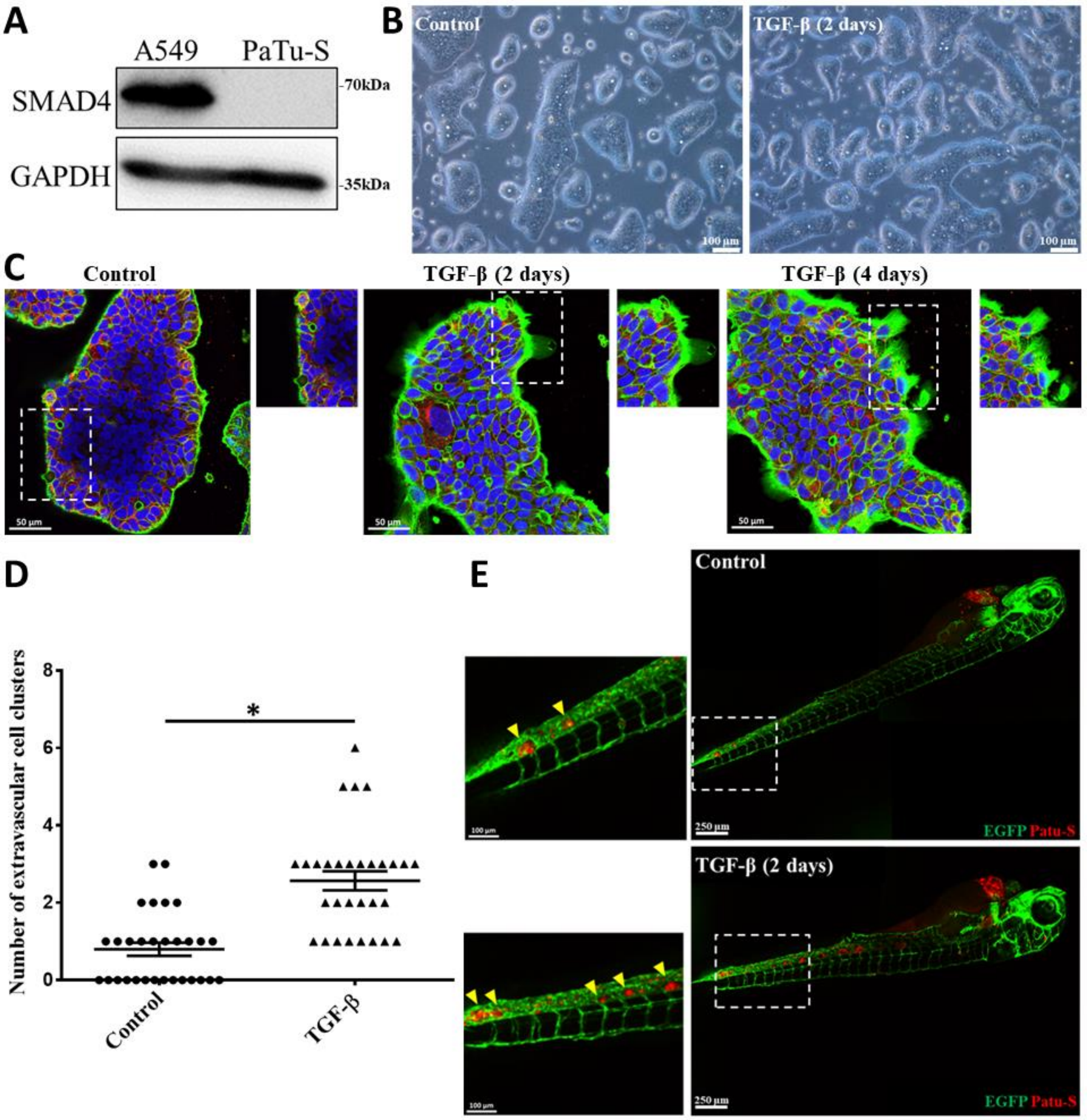


**Figure S1. Responses to TGF-β in PaTu-S cell line. (A)** The PaTu-S cell line was validated for its SMAD4 deficiency using Western Blot analysis by A549-VIM-RFP was included as a SMAD4 expressing cell line. **(B)** Morphological analysis of PaTu-S cells treated with vehicle control (Con) or TGF-β (2.5 ng/mL) for 2 days. Scale bar = 100 μm. **(C)** PaTu-S cells were double stained with anti-E-cadherin antibody and fluorescein-phalloidin to detect the expression of the epithelial marker, E-cadherin (red) and the formation of filamentous (F)-actin (green), respectively after treatment with vehicle control or TGF-β (2.5 ng/mL) for 2 days and 4 days. Nuclei are counterstained with DAPI (blue). Dashed boxes indicate the areas of the enlarged images that are shown in the right panels.

At borders of the cell colonies lamellipodia are visible; these structures were stimulated in response to TGF-β treatment. Images were captured with confocal microscopy. Scale bar = 50 μm. Experiments were performed in triplicate with similar results, and representative results are shown. **(D)** mCherry labeled PaTu-S cells were pretreated with vehicle control or TGF-β (2.5 ng/mL) for 2 days, then injected into zebrafish embryos. The number of extravasated cell clusters were analyzed at the fourth day after injection. Thirty embryos were analyzed in each group. *$P \leq 0.05$, unpaired Student's t test, n=1. **(E)** Representative images of zebrafish from control and TGF-β treatment groups with zoom in of extravasated cell on the left panel. Yellow arrows indicate the invasive cells. Scale bar = 250 μm or 100 μm.

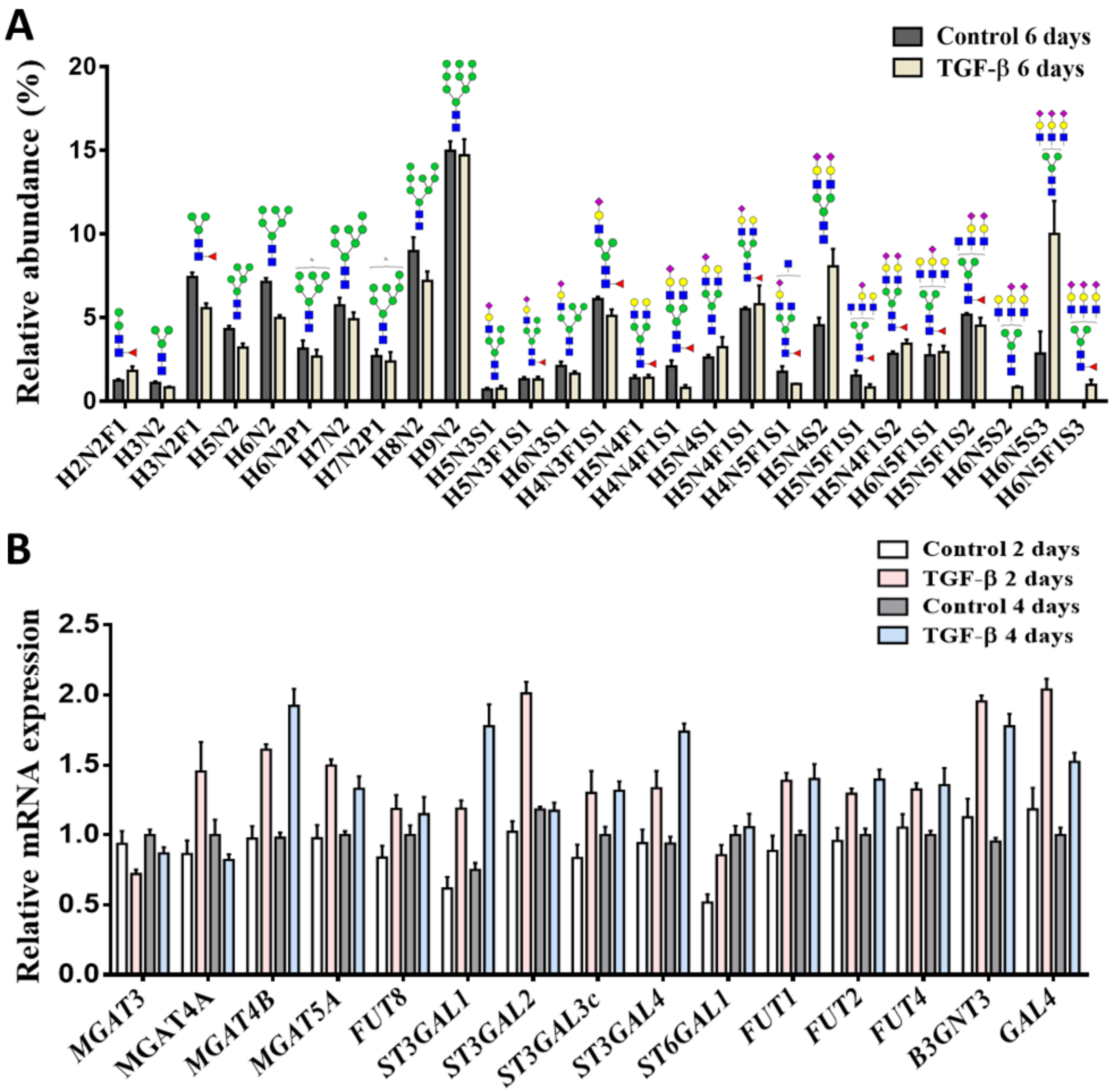


**Figure S2. Changes of *N*-glycosylation in PaTu-S cell line without or with TGF-β treatment. (A)** Relative quantification of individual *N*-glycans derived from 0.5 million PaTu-S cells treated with vehicle control or TGF-β (2.5 ng/mL) for 6 days as measured on PGC nano-LC-ESI-MS/MS in negative ion mode. Blue square: *N*-acetylglucosamine; yellow circle: galactose; green circle: mannose; red triangle: fucose; pink diamond: *N*-acetylneuraminic acid; P: phophate. **(B)** qRT-PCR analysis of *N*-glycosylation related gene expression levels in PaTu-S cells treated with vehicle control or TGF-β (2.5 ng/mL) for 2 days or 4 days. *GAPDH* mRNA levels were used for normalization. Fresh medium containing TGF-β or vehicle control was added every 2 days. Representative results are shown of three independent experiments or the data is expressed as the mean ± s.d. (n=3).

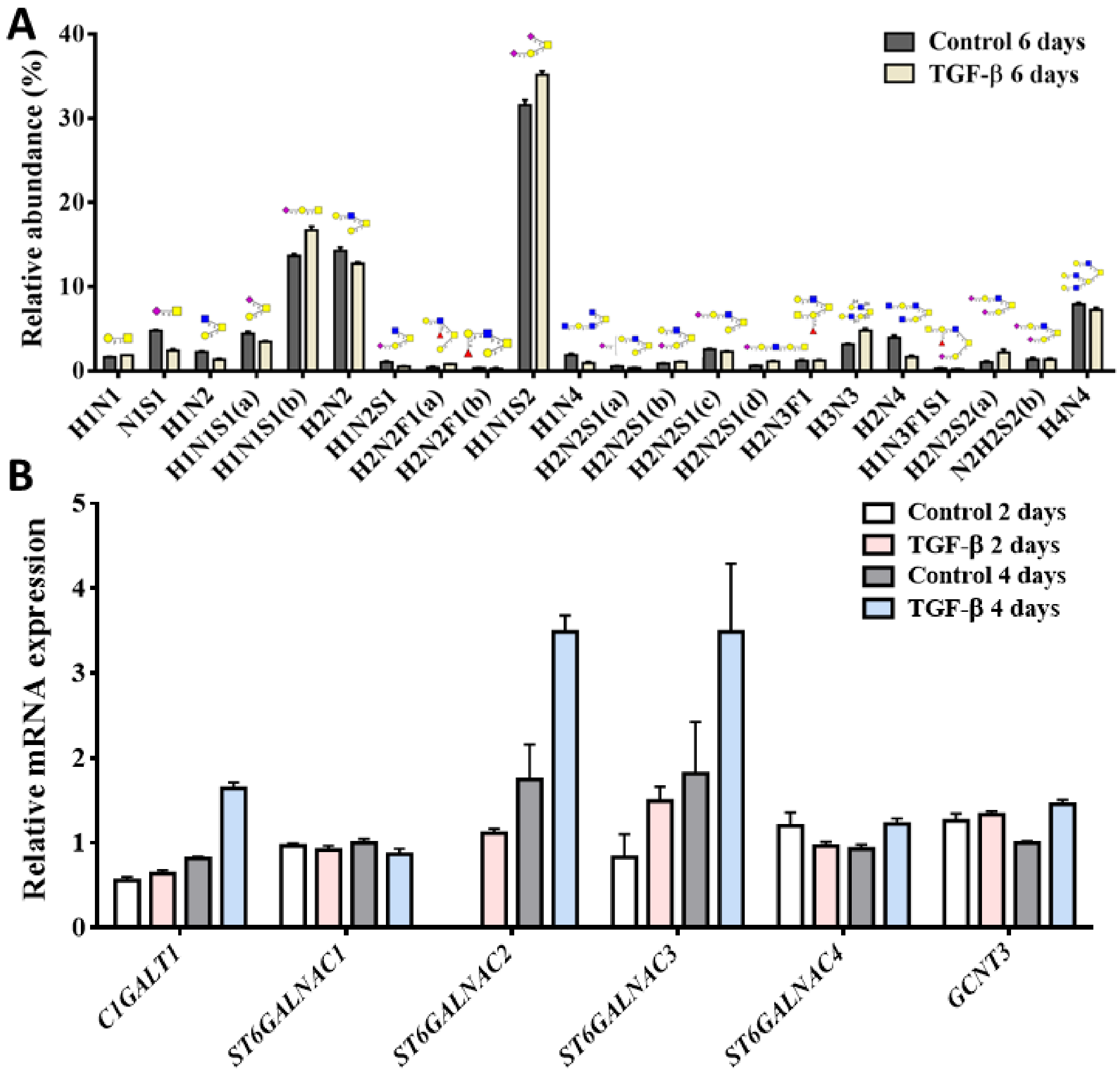


**Figure S3. Changes of *O*-glycosylation in PaTu-S cell line without or with TGF-β treatment.** **(A)** Relative quantification of individual *O*-glycans derived from 0.5 million PaTu-S cells treated with vehicle control or TGF-β (2.5 ng/mL) as measured on PGC nano-LC-ESI-MS/MS in negative ion mode. Blue square: *N*-acetylglucosamine; yellow circle: galactose; green circle: mannose; red triangle: fucose; pink diamond: *N*-acetylneuraminic acid. **(B)** qRT-PCR analysis of *O*-glycosylation related gene expression levels in PaTu-S cells treated with vehicle control or TGF-β (2.5 ng/mL) for 2 days or 4 days. GAPDH mRNA levels were used for normalization. Fresh medium containing TGF-β or vehicle control was added every 2 days. Representative results are shown of three independent experiments or the data is expressed as the mean ± s.d. (n=3).

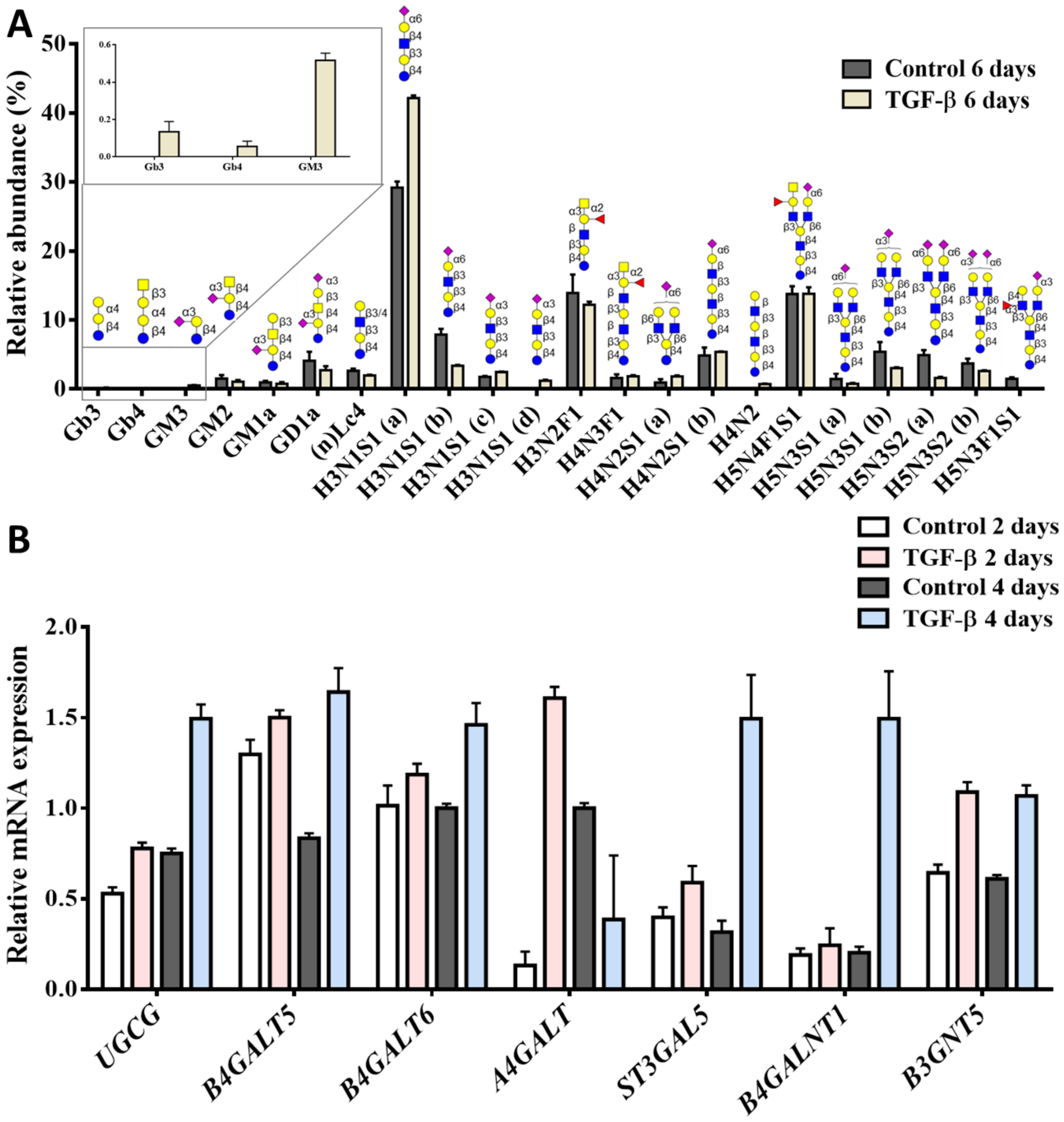


**Figure S4. Changes of glycosphingolipids (GSLs) in PaTu-S cell line without or with TGF-β treatment. (A)** Relative quantification of individual GSL-glycans derived from 0.5 million PaTu-S cells treated with vehicle control or with TGF-β (2.5 ng/mL) for 6 days as measured on PGC nano-LC-ESI-MS/MS in negative ion mode. Blue square: *N*-acetylglucosamine; yellow circle: galactose; green circle: mannose; red triangle: fucose; pink diamond: *N*-acetylneuraminic acid. **(B)** qRT-PCR analysis of GSL-related gene expression levels in PaTu-S cells treated with vehicle control or TGF-β (2.5 ng/mL) for 2 days or 4 days. GAPDH mRNA levels were used for normalization. Fresh TGF-β or vehicle control was added every 2 days. Representative results are shown of three independent experiments or the data is expressed as the mean ± s.d. (n=3).

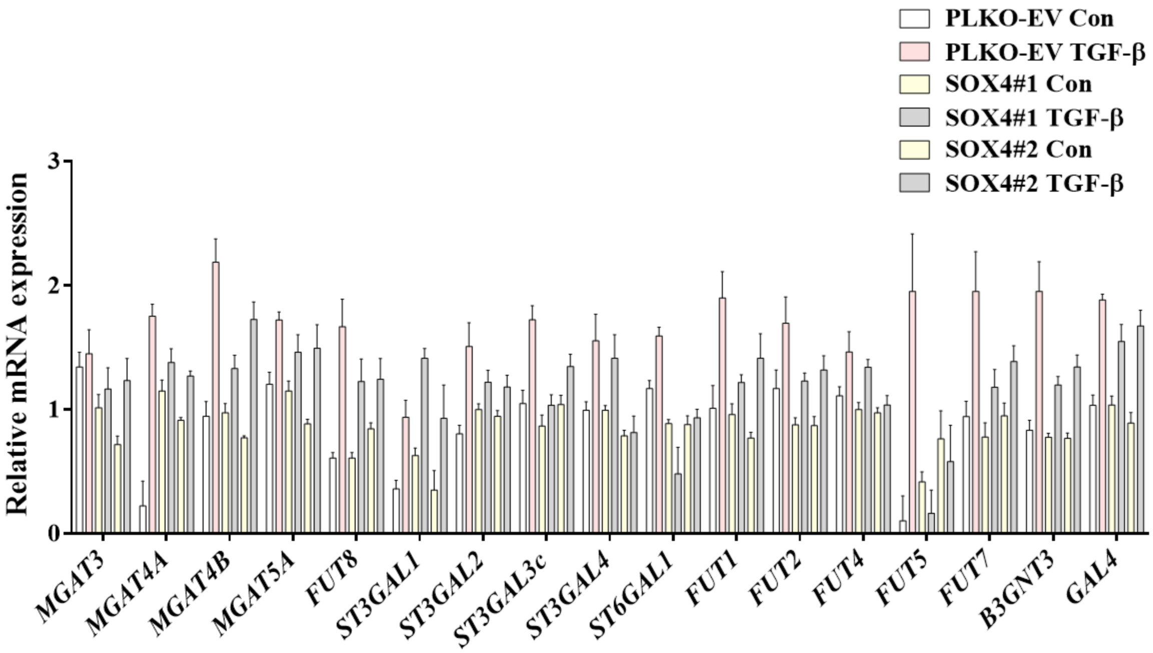


**Figure S5. mRNA expression levels of *N*-glycan biosynthesis related glycosyltransferases genes in PaTu-S cells with SOX4 deletion.** qRT-PCR analysis of *N*-glycosylation related genes in PaTu-S cells stably infected with two SOX4 shRNAs (sh#1 and sh#2) or empty vector shRNA (PLKO-EV) after treatment with vehicle control (Con) or TGF-β (2.5 ng/mL) for 2 days. *GAPDH* mRNA levels were used for normalization. Data are expressed as the mean ± s.d, of triplicates and are representative of at least two independent experiments.

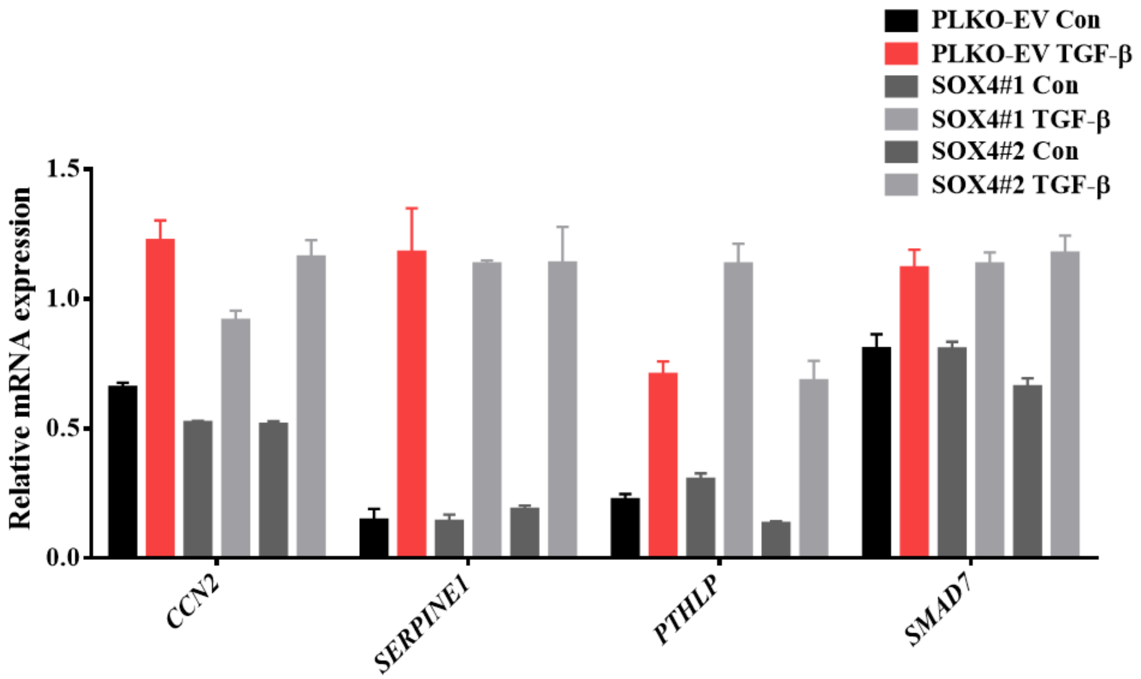


**Figure S6. mRNA expression levels of TGF-β target genes in PaTu-S cells with SOX4 deletion.** qRT-PCR analysis of *CCN2*, *SERPINE1*, *PTHLH* and *SMAD7* in PaTu-S cells infected with PLKO-EV, SOX4 sh#1 and SOX4 sh#2. after vehicle control (Con) or TGF-β (2.5 ng/mL) treatment for 6 h. *GAPDH* mRNA levels were used for normalization. Data are expressed as the mean ± s.d, of triplicates and are representative of at least two independent experiments.

**Table S1. Primers used in this study for quantitative real-time-polymerase chain reaction (qRT-PCR)**

| Species | Gene name | Forward (5' to 3') | Reverse (5' to 3') |
|---|---|---|---|
| Human | *GAPDH* | TGCACCACCAACTGCTTAGC | GGCATGGACTGTGGTCATGAG |
| Human | *CCN2* | TTGCGAAGCTGACCTGGAAGAGAA | AGCTCGGTATGTCTTCATGCTGGT |
| Human | *SERPINE1* | CACAAATCAGACGGCAGCACT | CATCGGGCGTGGTGAACTC |
| Human | *PTHLH* | AGACTGGTTCAGCAGTGGAG | TTCTTCCCAGGTGTCTTGAG |
| Human | *SMAD7* | TCCAGATGCTGTGCCTTCC | GTCCGAATTGAGCTGTCCG |
| Human | *CDH1* | CAGCCGCTTTCAGATTTTCAT | CCCGGTATCTTCCCCGC |
| Human | *CDH2* | CAGACCGACCCAAACAGCAAC | GCAGCAACAGTAAGGACAAACATC |
| Human | *SNAI2* | TGTTGCAGTGAGGGCAAGAA | GACCCTGGTTGCTTCAAGGA |
| Human | *VIM* | CCAAACTTTTCCTCCCTGAACC | CGTGATGCTGAGAAGTTTCGTTGA |
| Human | *MGAT3* | GCACTTCTTCAAGACCCTGTCCTA | AGAAAAAGCTGGACACCAGGTTA |
| Human | *MGAT4A* | CTGTGGAAGTTTTGCCTTTTAAGAG | TGAAATGGGATTGAGACTTGGA |
| Human | *MGAT4B* | CGGAGGACAAGCTCTTCAACA | AGGGCCTCCTTGTCTGACTGA |
| Human | *MGAT5A* | GATGTGCTTTTCTGAATCCCAAGT | GCCGCCCGATGAAAACT |
| Human | *FUT8* | TCTTCATCCCCGTCCTCCA | GAGACACCCACCACACTGCA |
| Human | *ST3GAL1* | GGGCAGACAGCAAAGGGAA | GGCCGTCACGTTAGACTCAAA |
| Human | *ST3GAL2* | ACAGGTGGACAGAGCATCAC | CCCGTACACGTTCACCTCAT |
| Human | *ST3GAL3c* | GGGTCACGAATTGACGACTATG | GTGATGCGCAGTGTCGTTTT |
| Human | *ST3GAL4* | ATAAGAAGCGGGTGCGAAAGGG | TCCGTGGCTGTTGCATTGGC |
| Human | *ST6GAL1* | CATCCAAGCGCAAGACTGACG | TGTGCCCTGGTTGAGATGCTTC |
| Human | *FUT1* | GCAGGCCATGGACTGGTT | CCTGGGAGGTGTCGATGTTT |
| Human | *FUT2* | CTCGCTACAGCTCCCTCATCTT | CGTGGGAGGTGTCAATGTTCT |
| Human | *FUT4* | GAGCTACGCTGTCCACATCACC | CAGCTGGCCAAGTTCCGTATG |
| Human | *FUT5* | GTCCCGAGACGATGCCACT | CCGGTGACAGGTTCCACTG |
| Human | *FUT7* | TCCGCGTGCGACTGTTC | ACCCTCAAGGTCCTCATAGACTTG |
| Human | *B3GNT3* | GTGGGACTTCCACGACTCCTT | GCACCTTGTCTCCTGCCACT |
| Human | *GAL4* | GTATAAGAGCCACCACCGCC | GTAGTAAGGCAGCGTCGGG |
| Human | *C1GALT1* | CCAGAGAAGCAAAGGTCACCA | TTCCCGAAAGTGTATTTCTGACATC |
| Human | *ST6GALNAC1* | GACGCTGACCCTTCGCTATT | TGCGTTGATGCACATAAGCC |
| Human | *ST6GALNAC2* | TCCGCGACTATGTGATGCTG | TGATTTCAAGAACCTGTCCCCT |
| Human | *ST6GALNAC3* | TGCCCAAATATACGTGACCACA | TCACTCTGTACTGTCCTTCCCA |
| Human | *ST6GALNAC4* | GTTCACCATGATCCTCGCG | TGACACTCATCTAGCCGGCC |
| Human | *GCNT3* | TGTTCACATTTGCTGCCACG | GTGGAGGAGGACACAATCCTTT |
| Human | *UGCG* | TGATCAGGTGGACCAAACTACG | ATCTGAACACATGGTGGGCT |
| Human | *B4GALT5* | TGCAGGCTATTCTGTGAGCC | CCTCAGCAGAGCATACCTTCC |
| Human | *B4GALT6* | CCATACCTCCCCTGTCCAGA | TTTTGGCCTCCAATGACCCC |
| Human | *A4GALT* | AACACTGGGGTGATGCAGG | AAATCCATCTCGATATTCTGCAGTG |
| Human | *ST3GAL5* | GCACCACTGTCTGACCTTGA | CCAGAATGGCAGGGTTTCCT |
| Human | *B4GALNT1* | TGAGGACCCTCAGGCCG | GCCACATCCTGTCTAACGCT |
| Human | *B3GNT5* | CCAGCGACTTTAGCTCCGAT | TTCCATGCCACCTCCAAGTC |
